# A desert endophyte, *Priestia megaterium* SI1-IITJ, improves fluoride stress tolerance by reducing fluoride content of plant tissues and perturbing salt tolerance and defense genes of *Arabidopsis thaliana*

**DOI:** 10.1101/2025.06.20.660123

**Authors:** Devanshu Verma, Pinki Sharma, Rishabh Kumar, Neelam Jangir, Vijesh Prajapat, Debankona Marik, Rabisankar Mandi, Trisikhi Raychoudhury, Nar Singh Chauhan, Ayan Sadhukhan

## Abstract

We isolated a fluoride (F^-^)-resistant bacterium, *Priestia megaterium* SI1-IITJ, from thse internal root tissues of several Thar Desert plants, *Aerva javanica*, *Cyperus conglomeratus*, *Senna tora*, and *Tephrosia purpurea*, tolerating up to 100 mM NaF. The root endophytic behavior of the isolate was confirmed by scanning electron microscopy. SI1-IITJ possesses plant growth-promoting properties, including auxin production (19.8 μg mL^-1^), phosphate solubilization (index 3.64), ACC deaminase (0.54 mmol α-ketobutyrate mL^-1^) and nitrate reductase (0.65 μmol mL^-1^ nitrite) activities, revealed by biochemical tests and whole genome sequencing. SI1-IITJ extrudes F^-^ from the cell, possibly through an F^-^ efflux transporter, *CrcB*, identified in its genome. Significant growth improvements were observed in *Arabidopsis thaliana* under F^-^ stress in hydroponics and soil culture upon coculture with SI1-IITJ, which improved the chlorophyll content by 1.6%, total nitrogen by 30.4%, and reduced reactive oxygen species by 48.9% and F^-^ content by 63.9% in plant tissues. A differential gene expression analysis of *A. thaliana* by transcriptome sequencing indicated an unperturbed F^-^ exporter, *AtFEX1,* but up-regulation of 55 genes regulating root meristem growth, cell wall modification, chlorophyll biosynthesis, Fe homeostasis, and high salt- and abiotic stress-responsive genes. On the other hand, 103 genes were down-regulated, suppressing systemic acquired resistance, plant defense, and H_2_O_2_ production. In conclusion, our results provide genomic insights into the mechanisms of F^-^ toxicity alleviation and plant growth enhancement by a desert PGPR, highlighting *Priestia megaterium* SI1-IITJ as a potential biofertilizer for mitigating F^-^ stress in plants.

## Introduction

Fluoride (F^-^) is a geogenic contaminant that is widely distributed in the environment, yet, unlike other pollutants, it is often overlooked despite being a serious threat to plant growth and overall soil health (Jha et al. 2009; Chatterjee et al. 2020; Mridha et al. 2021). At about 0.3 g kg^-1^, fluorine ranks as the thirteenth most prevalent element in the crust of the Earth and mostly occurs as sodium fluoride (NaF) (Bhat et al. 2015). It rarely exists in the elemental form, a highly reactive gas (Yadav et al. 2007). F^-^ naturally builds up in the soil through the weathering of parent rocks and minerals that contain F^-^, the deposition of volcanic emissions, the evaporation of surface waters, and the deposition of marine aerosols (Cronin et al. 2003; Lewandowska et al. 2013; Rafique et al. 2015). F^-^ occurs commonly in soil as cryolite (Na_3_AlF_6_), fluorite (CaF_2_), and apatite [Ca_5_(PO_4_)_3_F] (Mikkonen et al. 2018). Concentration of F^-^ in the soil ranges from 150 to 400 mg kg^-1^; however, in certain heavy clay-rich soils, concentrations exceeding 1000 mg kg^-1^ have been reported (Begum 2012). Plants absorb F^-^ through their roots and leaves, accumulating it within tissues at amounts that exceed those in the environment. Its accumulation usually depends on soil F^-^ levels, plant species, and soil properties (Kumar et al. 2021). Through a passive diffusive process, F^-^ enters the apoplast of roots. Although cell wall Ca^2+^ binds to F^-^, forming insoluble CaF_2_, and the cell membrane repels the negatively charged F^-^, HF is formed at the low apoplastic pH due to protonation of F^-^, which, being lipophilic, enters the cytoplasm through the plasma membrane easily. F^-^ easily passes the endodermal barrier, and through symplastic transport, enters the vascular tissues and eventually gets transported over long distances to all plant tissues. F^-^ induces toxicity in plants, decreasing crop yield and reducing food quality (Rizzu et al. 2021). F^-^ stress disrupts key processes such as chlorophyll production and photosynthesis, nutrient uptake, gene expression, and protein synthesis, reducing seed germination, plant growth, biomass accumulation, and productivity (Choudhary et al. 2019). Extensive exposure to F^-^ causes leaf chlorosis and necrosis, reduced root and shoot growth, and a decline in vigor index (Ghosh and Singh 2005; Baunthiyal and Ranghar 2014). F^-^ changes osmolyte levels, including proline and soluble carbohydrates, induces oxidative stress, and inhibits vital enzymatic activities (Sharma et al. 2012; Yadu et al. 2018; Rizzu et al. 2021).

Plant rhizospheres host plant growth-promoting rhizobacteria (PGPR), which enhance plants’ tolerance to various abiotic stresses and environmental pollutants (Khatoon et al. 2020). Hence, PGPRs can potentially substitute chemical fertilizers (Bhattacharya et al. 2020; Yang et al. 2022; Nie et al. 2023; Marik et al. 2024). Plant defense systems against pathogens and abiotic stress are activated by PGPRs, which release siderophores, organic acids, vitamins, and volatile chemical compounds (Chandran et al. 2021). Some bacterial species secrete ionophores that bind to anionic species. The bacteria can use or tolerate these complexes, which lowers the concentration of F^-^ in the environment, enabling F^-^ tolerance (Doble and Kumar 2005; Mukherjee et al. 2015, 2017). F^-^ interacts with ionophores associated with the microbial cell wall to diffuse into the cytoplasm, forming complexes with biological molecules. These complexes are then stored in the vacuole (Pal et al. 2022). By extruding F^-^, the FluC family of F^-^ efflux channels, encoded by the *crcB* genes, and the CLCF family of F^-^/H^+^ antiporters prevent this dangerous halide from building up in the bacterial cytoplasm (McIlwain et al. 2021). In eukaryotes, including fungi, plants, and some marine animals, F^-^ exporter (FEX) channels are involved in F^-^ tolerance (Baker et al. 2012; McIlwain et al. 2021). PGPRs isolated from high F^-^-containing soil were found earlier to increase plant biomass, root length, and chlorophyll content and reduce oxidative damage (Katiyar et al. 2024, 2025). But the precise molecular mechanisms of mitigation of F^-^ toxicity in plants triggered by F^-^ tolerant PGPRs remain elusive, prompting us to conduct this study.

The Thar Desert, spanning western India and parts of Pakistan, reports high F^-^ levels in soil and groundwater (Singh et al. 2012; Rafique et al. 2015; Thakur et al. 2023). The Thar is also a source of unique PGPRs, which help desert plants tolerate severe stress (Marik et al. 2024). Hence, we explored robust F^-^-resistant PGPR strains from several desert shrubs. We isolated a strain of *Priestia megaterium* that promoted plant growth and mitigated F^-^-induced toxicity in the plant model *Arabidopsis thaliana*. Key bacterial properties that encourage plant growth, such as auxin synthesis, phosphate solubilization, nitrate reductase, and ACC deaminase activities, were biochemically examined for the isolated bacterium. Its ability to tolerate F^-^ and colonize plant roots was also assessed. Whole genome sequencing was performed to identify bacterial genes associated with stress adaptation and plant interaction. The bacterial impact on plant growth and physiology was evaluated by measuring biomass, chlorophyll content, and the accumulation of F^-^ ions, nutrients, and reactive oxygen species (ROS) under F^-^ stress. A transcriptome analysis of bacteria-inoculated *A. thaliana* uncovered gene expression changes underlying the observed stress alleviation in plants. This comprehensive approach highlights *P. megaterium* as a promising biofertilizer improving plant health and resilience in F^-^-contaminated soils.

## Materials and Methods

### Fluoride-tolerant rhizobacteria isolation from Thar desert plants

Roots of several shrub species growing in the Indian Institute of Technology Jodhpur campus (26.4710° N, 73.1134° E), viz. *Aerva javanica*, *Tephrosia purpurea*, *Senna tora*, and a sedge, *Cyperus conglomeratus*, were exposed after digging the soil to a depth of 1 foot. Following their collection, the roots were subjected to surface sterilization using 70% ethanol, followed by treatment with 1% sodium hypochlorite and washes with sterile deionized water before aseptic maceration using phosphate-buffered saline (PBS), pH 7.4. An NaF stock solution was filter-sterilized, and the requisite amount was added after autoclaving the Luria Bertani (LB) agar before solidification. Serial dilutions of crushed roots in PBS were spread on the NaF-infused LB plates. The plates were left to grow overnight at 37°C. The resultant bacterial colonies were repeatedly streaked on LB agar plates containing 100‒200 mM NaF to obtain pure bacterial cultures. Gram staining and light microscopy confirmed the purity of the isolates.

### DNA extraction and de-duplication of isolates

The HiPura Bacterial Genomic DNA Purification Kit (Himedia, Thane, India) was employed to isolate bacterial genomic DNA. A PCR was conducted from genomic DNA using 16S universal primers (Table S1) using an Agilent SureCycler 8800 (Agilent Technologies, Mumbai, India). The Wizard^®^ SV Gel and PCR Clean-Up Kit (Promega, New Delhi, India) was used to clean the products. The fragments were digested by a four-base cutter HaeIII (SRL, Mumbai, India). The digested fragments were resolved on a 2% agarose gel. Bacterial DNA resulting in a similar band pattern was considered a duplicate.

### Assessment of root colonization of fluoride-tolerant rhizobacteria

Seeds of a host plant, *Senna tora*, were germinated in deionized water and transferred to ¼-Hoagland’s medium, pH 5.8 (Hoagland and Arnon 1938). The seedlings were grown under hydroponic conditions in the presence or absence of the bacterial isolate. After the growth period, roots were gently rinsed with PBS to remove loose surface-associated bacteria. This was followed by sample fixation for electron microscopic observation of tightly bound surface bacteria. For observation of bacteria colonizing the internal tissues, first, the root surfaces were sterilized with 70% ethanol and 1% NaOCl for 10 min and washed in PBS. Transverse sections of the root samples were prepared using a sterile surgical blade and placed on pieces of glass slides inserted in separate wells of 24-well plates. Subsequently, 2.5% glutaraldehyde was added to the wells. Following 1 h fixation, the samples underwent ten minutes of repeated washing in 0.1 M phosphate buffer. After five minutes of dehydration using 30%, 50%, 70%, 90%, 95%, and 100% ethanol, the samples were further dried for one hour in a vacuum desiccator. Carbon adhesive tapes were used to paste samples on aluminum stubs. Finally, silver sputter-coating using a Magnetron DC sputtering System (Scientific & Analytical Instruments, New Delhi, India) was conducted before imaging with an EVO18 special edition scanning electron microscope (Carl Zeiss, New Delhi, India).

For the colony count assay for bacterial colonization, *A. thaliana* seedlings were grown hydroponically for 14 days under sterile conditions, either in the presence or absence of the bacterial isolate. After the growth period, roots were carefully washed with sterile PBS to remove loosely attached, non-colonizing bacterial cells. The roots were then excised using a sterile surgical blade and weighed. To assess surface colonization, the root samples were rinsed three times with PBS, followed by vortexing for 30 s to dislodge the surface-associated bacteria. For internal tissue colonization analysis, root surfaces were sterilized with 70% ethanol and 1% NaOCl for 10 min and then rinsed in sterile PBS repeatedly. The sterilized roots were then mechanically crushed using a sterile mortar and pestle and resuspended in PBS. Both surface and internal root colonization preparations were spread and allowed to grow on chloramphenicol (25 µg mL^-1^) or ampicillin (100 µg mL^-1^) plates at 37°C before counting colonies.

### Biochemical characterization of fluoride-tolerant rhizobacteria

To test the F^-^ tolerance of the rhizobacterial isolate, 50 µL of an overnight-grown culture in liquid LB medium (OD_600 nm_ = 1), was used to inoculate fresh LB media supplemented with 0‒200 mM NaF, and the OD_600 nm_ was recorded at time intervals. The isolate was cultured in LB medium with 100 mM NaF for further biochemical characterization. For the catalase test, 10 µL of culture was spotted on a clean glass slide, followed by 3% hydrogen peroxide (H_2_O_2_) to observe effervescence (Reiner 2010). Oxidase activity was evaluated using a colorimetric method by mixing 0.3 mL of 1% 4-aminodimethylaniline oxalate and 0.2 mL of 1% 1-naphthol with 4.5 mL of culture and shaking vigorously. A blue coloration within a few minutes indicated a positive oxidase reaction (Shields and Cathcart 2010). For the starch hydrolysis test, starch agar medium was prepared using 10 g L^-1^ soluble starch, 3 g L^-1^ beef extract, and 12 g L^-1^ agar, pH 7.5. Bacteria were inoculated on the medium and grown at 37°C for 48 h. Thereafter, iodine solution was added to the plates to detect starch degradation (Lal and Cheeptham 2012). Bacterial strains were inoculated in methyl red–Voges Proskauer (MR–VP) broth, composed of 5 g L^-1^ dextrose, 7 g L^-1^ peptone, and 5 g L^-1^ K_2_HPO_4_, and incubated at 37°C for 24 h. For the MR test, a drop of methyl red was added to 1 mL of the culture (McDevitt 2009). For the VP test, 1 mL of the same culture was mixed with 0.6 mL of 5% 1-naphthol and 0.2 mL of 40% KOH, incubated at 37°C for 30 min, and the color was observed (McDevitt 2009). Citrate utilization was assessed using Simmons’ citrate agar medium containing 2 g L^-1^ sodium citrate, 5 g L^-1^ NaCl, 0.08 g L^-1^ bromothymol blue, 1 g L^-1^ K_2_HPO_4_, 1 g L^-1^ NH_4_H_2_PO_4_, 0.2 g L^-1^ MgSO_4_·7H₂O, and 15 g L^-1^ agar, pH 6.9. Bacteria were grown on the surface of citrate agar slants at 37°C for 24 h. A green-to-blue color change of the medium due to alkalinization indicated citrate utilization (MacWilliams 2009).

### Assessment of plant growth-promoting traits of bacteria

#### Assessment of auxin production

The ability to synthesize auxin indole acetic acid (IAA) was analyzed by growing bacterial cultures at 37 °C for 24 h in liquid LB medium with 0.2% L-tryptophan. First, the bacterial cells were separated by centrifugation and the cell-free broth incubated for 15 min with Salkowski’s reagent, composed of 0.5 M FeCl_3_ and 35% HClO_4_ in the ratio 1:50. Red coloration indicated IAA formation, quantified spectrophotometrically at 530 nm against an IAA standard curve (R^2^ = 0.979) (Bric et al. 1991).

#### Assessment of phosphate solubilization

To evaluate the phosphate-solubilizing activity of the bacterial isolates, overnight-grown cultures were resuspended in PBS, adjusting their optical density (OD) to 1.0, corresponding to approximately 8 × 10⁸ cells mL^-1^. Subsequently, 1 µL of this bacterial suspension was spot-inoculated onto a phosphate solubilization medium, composed of 0.5 g L^-1^ yeast extract, 5 g L^-1^ Ca_3_(PO_4_)_2_, 0.2 g L^-1^ (NH_4_)_2_SO_4_, 1 g L^-1^ MgSO_4_·7H_2_O, 0.1 mg L^-1^ MnSO₄, 0.1 mg L^-1^ FeSO_4_, 10 g L^-1^ glucose, and 20 g L^-1^ agar, adjusted to pH 7 (Pikovskaya 1948). The plates were incubated at 37 °C. After a 7-day growth, the colony diameter (CD) and the halo zone diameter (HZD) surrounding the colony were measured. The solubilization index (SI) was calculated using the formula SI = (HZD + CD) / CD.

For the quantitative analysis of phosphate solubilization, bacterial cultures were grown for 7 days at 37 °C in a phosphate growth medium. This medium contained either 5 g L^-1^ Ca_3_(PO_4_)_2_ or CaHPO_4_ as the insoluble phosphate source, along with 5 g L^-1^ MgCl_2_.6H₂O, 5 g L^-1^ MgSO_4_.7H₂O, 0.2 g L^-1^ KCl, 0.1 g L^-1^ (NH_4_)_2_SO_4_ and 10 g L^-1^ glucose (Mehta and Nautiyal 2001; Sharma et al. 2007). After growth, the culture was centrifuged at 5,000 rpm for 10 min. Next, 0.4 mL HClO_4_, 0.2 mL of 0.25% 1-amino-2-naphthol-4-sulfonic acid (prepared in a solution of 195 mL 15% NaHSO_3_ and 5 mL 20% Na_2_SO_3_), and 0.4 mL of 2.5% (NH_4_)_6_Mo_7_O_24_ (in 5 N H₂SO₄) were added to 5 mL of the supernatant. The absorbance was measured at 640 nm (A_640_) and quantified against a standard curve (R^2^ = 0.9877) prepared with serial dilutions of KH_2_PO_4_ (Fiske and Subbarow 1925).

#### Assessment of nitrate reductase activity

After inoculating a single microbial colony in 5 mL of sterile LB broth (pH 7.0) with 0.1% potassium nitrate, the culture was grown at 37°C until an OD_600 nm_ of 1.0. It was then centrifuged for 5 min at 16,000 g. Nitrite production was calculated using the protocol described by Streeter and Devine (1983). One mL of 0.02% N-(1-naphthyl) ethylenediamine and 1 mL of 1% sulfanilamide were added to 1 mL of the bacterial supernatant. The color change from colorless to pink was observed, and the A_550_ was measured. Nitrite produced in the reaction was quantified against a standard curve (R^2^ = 0.99827) prepared with different NaNO_2_ concentrations. All assays were performed in triplicate.

#### Assessment of ACC deaminase activity

The Maheshwari et al. (2020) procedure was used to evaluate the microbial isolate’s production of aminocyclopropane-1-carboxylic acid (ACC) deaminase. Microbial cells were cultured in minimal media (Dworkin and Foster 1958) and harvested by 5 min centrifugation at 16,000 g. By evaluating the amount of α-ketobutyrate in the reaction mixture against a standard curve (R^2^ = 0.9998) generated from various α-ketobutyrate concentrations, the ACC deaminase activity was determined. All assays were conducted in triplicate.

#### Whole genome sequencing of fluoride-tolerant rhizobacterial isolates

De-duplicated bacteria were subjected to whole-genome sequencing on a Nanopore MinION MK1C platform utilizing the Midnight pipeline. The Unicycler assembler, which incorporates SPAdes as its *de novo* assembler, was used to put together the resultant lengthy reads (Wick et al. 2017). The assembled genome was annotated using Prokka version 1.12. The 16S rRNA gene sequence was extracted from the whole genome sequence, and a nucleotide BLAST search against the non-redundant database was performed to identify homologous sequences. Based on the highest percentage of sequence identity, a taxonomic classification was assigned to the isolate using MEGA-X software, using the default parameters. For comparative analysis, the genomes of several *Priestia* species were downloaded from NCBI. Phylogenomic relationships between the isolate and reference *Priestia* sp. genomes were inferred using the Roary platform ver. 3.11.2. A circular genome map was drawn using the Proksee tool, providing a graphical representation of the genomic structure and annotated features (Grant et al. 2023). Phylogenomic analysis was performed using FastTree ver. 2.1.10. The Comprehensive Antibiotic Resistance Database (CARD) identification was used to predict the bacterial genes conferring antibiotic resistance (McArthur et al. 2013). The mobile genetic elements or mobilome were predicted by VRprofile2 (Wang et al. 2022). The genes for F^-^ stress resistance and plant growth promotion were also identified from the genome.

#### Assessment of plant growth and fluoride tolerance promotion of rhizobacteria

Seeds of *A. thaliana*, ecotypes Col-0 and Star-8, were surface sterilized using 70% ethanol and 2% NaOCl and washed several times with sterile deionized water. Control seeds were then immersed in sterile PBS, whereas the treatment seeds were soaked in PBS containing suspended bacteria (OD_600 nm_ = 1). Then the seeds were stratified at 4°C for three days. The imbibed seeds were sown on nylon meshes with a pore size of 50 μm. The meshes were mounted on photographic frames and floated on 300 mL ¼-Hoagland’s solution, pH 5.8, in sterile magenta boxes with partially open lids. The sides of the magenta boxes were covered with aluminum foil, preventing light exposure to the roots. The ¼-Hoagland’s solution was supplemented with 1 mM NaF for F^-^ treatment before adjusting the pH to 5.8 with HCl/NaOH. The boxes were incubated at 22 ± 3°C, 60 ± 5% humidity, and 12 h photoperiod supplied through T8 LED lamps emitting 80 μmol m^-2^s^-1^ of photosynthetically active radiation. After sowing seeds on the meshes, the solution was inoculated with 1 mL of PBS-resuspended bacteria (OD_600 nm_ = 1), or sterile PBS for controls. The media inoculation was done twice at an interval of seven days. The roots could grow downwards into the solution upon germination, and shoots emerge upwards from the meshes. After two weeks, the plants were carefully picked from the meshes with forceps, placed on agar, and photographed from an equal distance for measuring the root and shoot dimensions using the LIA 3.2 software, and average values of ten plants were presented with standard error (Marik et al. 2024).

For the soil culture experiment, *A. thaliana* was grown similarly in hydroponics for two weeks. Thereafter, uniformly sized plants were picked up from the meshes using forceps and planted in 2-inch pots containing equal amounts of autoclaved soil mix composed of 2:1:1 soil rite (Keltech Energies Ltd., Bangalore, India): desert soil: vermiculite. After 2 days of plant acclimatization in soil, some pots were inoculated with the overnight-grown bacterial isolate suspended in PBS (OD_600 nm_ = 1), and others with sterile PBS for the ‘no-bacteria’ controls. Inoculation was repeated three times at an interval of 7 days. After 10 days, the pots were irrigated thrice with 0, 100, or 150 mM NaF solutions at an interval of 7 days. Every 3^rd^ day, all plants were watered with deionized water. Finally, after 30 days of growth in the soil, the plants were photographed from the top with a scale. Rosettes were excised, washed with deionized water, blotted with tissue paper, and weighed. Roots were separated carefully from the soil by being dipped in water and gently shaken. The roots were washed a few times, blotted, and weighed to determine the biomass. The average values of ten plants were presented with standard error.

#### Transcriptome and real-time PCR analysis

*A. thaliana* ecotype Col-0 was cocultured in hydroponics with SI1-IITJ under 1 mM NaF as described in section 2.7. After two weeks, non-inoculated control and bacteria-inoculated seedlings were washed in deionized water and harvested by snap freezing using liquid nitrogen for RNA isolation using the CTAB-LiCl method. Three biological replicates for each of the control and bacteria-inoculated samples were used, each replicate consisting of five seedlings. Agilent 4150 TapeStation system and Qubit 4 fluorometer (Thermo Fisher Scientific, Mumbai, India) ascertained the quality of the RNA before its conversion into cDNA libraries using NEB NebNext Ultra II RNA library kit. Illumina NovaSeq 6000 V1.5 platform was used to sequence quality-passed libraries in 2 × 150 bp paired-end reads. AdapterRemoval ver. 2.3.2 (Schubert et al. 2016) was used to trim adapters from the raw sequence reads. Hisat2 ver. 2.2.1 (Zhang et al. 2021) was used to align the reads to the *A. thaliana* genome (GCF000001735.4). FeatureCounts v2.0.3 (Liao et al. 2013) was employed to determine the gene expression levels, and edgeR (Chen et al. 2025) was used to determine the differentially expressed genes (DEGs). To learn more about DEGs’ roles in particular biological processes, they were subjected to a Gene Ontology (GO) functional enrichment analysis using g:Profiler (Kolberg et al. 2023). For real-time quantitative PCR (qPCR) validation of the transcriptome results, the total RNA was converted into cDNA using RevertAid RT reverse transcriptase kit (Thermo) and used as a template. Primers were designed using the Primer3 tool (Kõressaar et al. 2018). *AtUBQ1* was used as an internal control.

#### Network analysis of differentially expressed genes

Protein–protein interaction (PPI) networks of the up- and down-regulated genes were visualized using STRING (Szklarczyk et al. 2022). In both networks, node size represents the degree of connectivity, and a color scale indicates the gene expression pattern. Colored rings around the nodes denote their association with significantly enriched biological pathways. For hub gene identification, a more stringent interaction confidence score of 0.7 (high confidence) was applied in the STRING database to ensure the inclusion of only highly reliable interactions. The resulting high-confidence network was subsequently imported into Cytoscape, where key hub genes were identified using cytoHubba (Ono et al. 2025). Hub gene analysis was performed using topological analysis algorithms, specifically the Maximal Clique Centrality (MCC) method. Additionally, MCODE analysis was conducted to detect densely connected modules within the network, which were further refined to final hub genes based on their MCC scores, biological relevance, and having a node degree ≥ 2. These hub genes were prioritized for their potential key roles in the underlying biological processes.

#### Assessment of plant chlorophyll, nutrient, and ROS content

For chlorophyll measurements, 0.5‒1.0 g of fresh shoot tissue from soil-grown plants (section 2.7) was macerated in 20 mL of 80% acetone and centrifuged at 14,000 rpm for 15 min. A_663_ and A_645_ of the supernatant were measured using a UV-1800 UV-visible spectrophotometer (Shimadzu, New Delhi, India). The following formula was used for estimation of concentration from absorbance values: chlorophyll a (g L^-1^) = 0.0127A_663_ – 0.00259A_645_, chlorophyll b (g L^-1^) = 0.0229A_645_ – 0.00468A_663_, and total chlorophyll (g L^-1^) = 0.00802A_663_ + 0.0202A_645_. Finally, the chlorophyll content was normalized by fresh weight of tissue (mg g^-1^) (Arnon 1949).

For total tissue nitrogen estimation, shoot tissues were homogenized in deionized water using a mortar and pestle, and the homogenate was centrifuged at 14000 rpm for 5 min to settle the tissue debris. After transferring the supernatant into fresh tubes, 2 mL was combined with 5 mL of the oxidizing agent for each assay. Three mL of TN reaction mixture was then added and incubated for 30 min at 45°C before recording A_545_. The oxidizing agent was prepared by mixing 6 g L^-1^ H_3_BO_3_ with 10 g L^-1^ K_2_S_2_O_8_ in 0.075 M NaOH. The TN mix was composed of 200 mL of 0.54% VCl_3_ in 1.2 N HCl, 40 mL of 1% sulphanilamide in 2.4 N HCl, and 40 mL of 0.07% N-(1-naphthyl)-ethylenediamine (Pradhan and Pokhrel 2013). A_545_ values were converted to mmol nitrogen using a standard curve of Na_2_EDTA (R^2^ = 0.981).

To estimate total phosphate content, *A. thaliana* shoots were homogenized in deionized water and centrifuged at 14,000 rpm for 15 min. Next, 0.5 mL of 10 N H_2_SO_4_, 2 mL of 2.5% (NH_4_)_6_Mo_7_O_24_, and 1 mL of 0.5 M hydrazine were added sequentially. The final volume was raised to 25 mL with water and incubated at 25°C for 45 min. A_840_ was recorded, and the concentration was estimated using a standard curve of KH_2_PO_4_ (R^2^ = 0.9) (Hartmann and Asch 2018).

The amount of H_2_O_2_ inside tissues was determined using the Elstner and Heupel (1976) method. After homogenization of 1 g of fresh shoot tissue in 1 mL of 5% HClO_4_, centrifugation for 15 min at 12,000 rpm, 0.4 mL of 10 mM (NH_4_)_2_Fe(SO_4_)_2_·6H_2_O, 0.2 mL of 2.5 M KSCN, and 0.4 mL of 50% C_2_HCl_3_O_2_ were added to the supernatant. After mixing carefully, A_480_ was recorded. A_480_ values were converted to mM H_2_O_2_ using a standard curve of pure H_2_O_2_ dilution series (R^2^ = 0.9989) (Sagisaka 1976). All values were represented per gram of tissue fresh weight.

#### Assessment of fluoride content in bacteria and plant tissues

To measure the F^-^-content inside the bacterial cells and the culture broth outside, after 24 h of growth, cells were centrifuged at 5,000 rpm for 15 min. A total ionic strength adjustment buffer (TISAB) was mixed with the supernatant in a 1:1 ratio after it had been diluted 1000 times with PBS. TISAB contained 5.7% v/v of glacial acetic acid, 57 g L^-1^ of NaCl, 2 g L^-1^ of EDTA, and 7 g L^-1^ of sodium citrate, pH 5.3 (Katiyar et al. 2024; Prajapat and Raychoudhury 2025). An Orion star A214 F^-^ ion selective electrode (Thermo Fisher) was used to measure the F^-^ concentration. The probe was calibrated with a dilution series of NaF before measurement. The pellet was resuspended in 10 mL of PBS, sonicated using a PRO656 probe sonicator (Labman Scientific Instruments Pvt. Ltd., Chennai, India) for 40 min at 20% power ratio, placing the tube on ice, and similarly mixed with 1:1 TISAB before measurement. To determine the F^-^ content inside plant tissues, rosettes were harvested after the soil culture experiment and weighed. The samples were dried for 2 days at 70°C, ground into powder, and suspended in 2.5 mL of 0.1 N HClO_4_ in a 15 mL polypropylene tube. Thereafter, TISAB was mixed with the samples and measured using the F^-^ ion selective electrode. F^-^ concentration was expressed as μmole g^-1^ tissue fresh weight.

### Statistical analysis

All growth and biochemical experiments were repeated three times. To rule out environmental effects, the growth experiments in pots followed a randomized complete block design, consisting of 10 plants per treatment. All biochemical tests consisted of three independent biological replicates for each treatment. Two samples were compared using Student’s *t*-test. The transcriptome results were statistically treated with Fisher’s exact test after Benjamini-Hochberg *false discovery rate* (*FDR*) correction at *P* < 0.05.

## Results

### Thar desert plants contain fluoride-tolerant endophytic bacteria having plant growth-promoting properties

We isolated five pure cultures of F^-^-tolerant bacteria from the internal root tissues of the Thar Desert shrubs *Aerva javanica*, *Tephrosia purpurea*, *Senna tora*, and a sedge, *Cyperus conglomeratus* (Fig. 1A), selected on LB media with 100 mM NaF. Each isolate was a Gram-positive rod, named SI1-IITJ, SI2-IITJ, GI4-IITJ, GI6-IITJ, and BI3-IITJ (Fig. 1B). A restriction fragment analysis using the four-base-cutter HaeIII on the amplified 16S rRNA gene fragment indicated that all bacteria were alike (Fig. 1C). This was verified using whole-genome sequencing analysis. Hence, we report all further analyses with one representative isolate, SI1-IITJ.

**Fig. 1.**
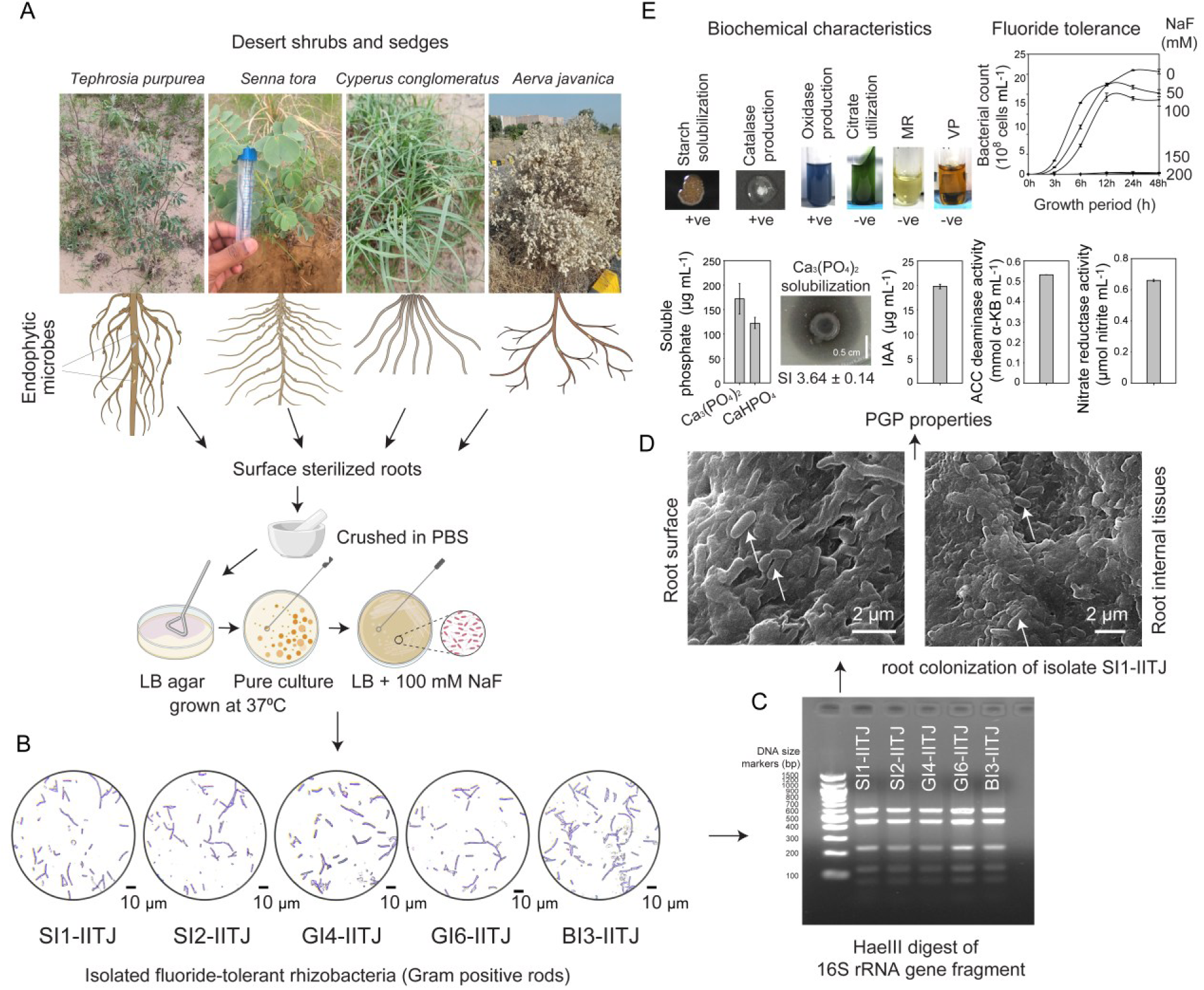
Isolation and characterization of fluoride-tolerant bacteria from the roots of desert plants. **(A)** Flow diagram of isolation of fluoride (F^-^)-tolerant bacteria from root internal tissues of desert plants: *Tephrosia purpurea*, *Senna tora*, *Cyperus conglomeratus*, and *Aerva javanica*. **(B)** Gram staining and light microscopic images of five pure F^-^-tolerant bacterial isolates, SI1-IITJ, SI2-IITJ, GI4-IITJ, GI6-IITJ, and BI3-IITJ. **(C)** De-duplication analysis by restriction digestion of an amplified fragment of the 16S rRNA gene using a four-base cutter HaeIII. The banding pattern indicates the same bacteria in all the samples **(D)** Scanning electron micrographs (SEM) of the root surface and root transverse section of a host plant *Senna tora,* with rod-shaped bacteria indicated by arrows. The plants were grown from seeds and inoculated with a pure culture SI1-IITJ. After 15-day growth, roots were harvested and processed for SEM (see Methods). The white bars indicate the scale in micrometers. **(E)** Biochemical characteristics, F^-^ tolerance, and plant growth-promoting traits of the isolate SI1-IITJ. The results of starch solubilization, catalase and oxidase production, citrate utilization, methyl red (MR), and Voges-Proskauer (VP) tests with SI1-IITJ are shown. Growth curves of bacteria in liquid LB supplemented with 0, 50, 100, and 150 mM NaF are demonstrated throughout 48 h. Other plant growth-promoting characteristics, including phosphate solubilization, indole acetic acid (IAA) production, 1-aminocyclopropane-1-carboxylate (ACC) deaminase, and nitrate reductase activities are also shown (see Methods). Average values of three replicates are presented with standard error.

We tested the ability of SI1-IITJ to colonize the roots of its host plant, *Senna tora,* and the model plant *A. thaliana*. Scanning electron microscopy indicated bacterial colonization on both the root surface and internal tissues inside root cross sections (Fig. 1D). We had tested the growth of SI1-IITJ in different antibiotics: kanamycin, gentamicin, chloramphenicol, and ampicillin, to find that this bacterium could tolerate 100 mg mL^-1^ ampicillin and 50 mg mL^-1^ chloramphenicol, producing colonies on overnight incubation. The colony count assay revealed that in *A. thaliana* root surfaces, bacterial counts of 1.64 × 10^6^ ± 0.8 × 10^6^ CFU g^-1^ root FW and in root internal tissues, 4.2 × 10^5^ ± 1.2 × 10^5^ CFU g^-1^ root FW were observed in ampicillin plates, whereas 1.21 × 10^6^ ± 0.5 × 10^6^ g^-1^ root FW and 2.65 × 10^4^ ± 0.4 × 10^4^ CFU g^-1^ root FW were observed in the root surface and internal tissues, respectively, in chloramphenicol plates. This indicates a host-independent root endophytic behavior of SI1-IITJ.

SI1-IITJ could tolerate an NaF concentration up to 100 mM in liquid LB media, while concentrations of 150 mM and 200 mM inhibited its growth (Fig. 1E). After a 24 h growth in liquid LB with 100 mM NaF, 96.9 ± 3.1 mM F^-^ remained in the supernatant, while 1.3 ± 0.1 mM F^-^ was detected in the pellet (Fig. S1). This suggests the bacterium extrudes F^-^ out of its cell to survive. Regarding other biochemical characteristics, SI1-IITJ showed a positive reaction due to the immediate formation of effervescence (oxygen bubbles) in the catalase test. The isolate exhibited a distinct blue color, confirming oxidase production. *Escherichia coli* was used as a negative control, and no color change was observed. Halos around the bacterial colonies indicated positive starch hydrolysis, demonstrating the ability of the strain to secrete amylase and utilize starch as a carbon source. *E. coli* served as a negative control and produced no clear zone. SI1-IITJ did not develop any red coloration, indicating a negative MR reaction. *E. coli* was used as a positive control and exhibited a distinct red color, confirming the validity of the test. No color change was observed in SI1-IITJ, indicating a negative VP reaction. *Enterobacter cloacae* was used as a positive control, and the characteristic pink-red color was developed (Marik et al. 2024). SI1-IITJ did not produce a color change, indicating a negative citrate utilization test result, while *Enterobacter cloacae*, used as a positive control, showed a distinct blue coloration (Fig. 1E).

SI1-IITJ produced 19.8 ± 0.13 μg mL^-1^ of IAA in liquid LB media supplemented with tryptophan, indicating its potential root growth enhancement properties (Fig. 1E). In solid media with insoluble Ca_3_(PO_4_)_2_, SI1-IITJ produced halo zones around its colonies with a solubilization index of 3.64 ± 0.14. SI1-IITJ could produce 171.9 ± 31.2 μg mL^-1^ and 121.7 ± 33.6 μg mL^-1^ of soluble phosphate when grown in liquid NBRI media containing two insoluble/ sparingly soluble phosphate sources, Ca_3_(PO_4_)_2_ and CaHPO_4_, respectively (Fig. 1E). Media without bacterial inoculation, processed similarly, showed no detectable soluble phosphate in the supernatant. SI1-IITJ exhibited nitrate reductase activity. The significant rise in nitrite levels (0.65 ± 0.01 μmol mL^-1^) in the reaction suggests that the microbial isolate effectively stimulates the activity of nitrate reductase, the enzyme converting nitrate to nitrite. SI1-IITJ was found to produce ACC deaminase (0.54 ± 0.005 mmol α-ketobutyrate mL^-1^), indicating its potential involvement in the removal of stress-induced ethylene in plants (Fig. 1E). However, we failed to detect any siderophore or exopolysaccharide-producing ability in SI1-IITJ.

### The genome of SI1-IITJ contains genes for fluoride tolerance and plant growth promotion

The assembly of the sequenced dataset of SI1-IITJ resulted in the generation of 33 contigs, accounting for a total of 6,242,625 bp in size, with 37.09% GC content and an N50 length of 430,874. Functional annotation of assembled contigs revealed one circular chromosome with 6,395 coding sequences, 46 rRNAs, 125 tRNAs, and one tmRNA. The 16S rRNA gene of SI1-IITJ exhibited 99.48% sequence identity with *Priestia megaterium* strain ATCC 14581, confirming its identification as a strain of *Priestia megaterium*. This conclusion was further supported by the phylogenetic analysis using the 16S rRNA gene (Fig. 2A). Based on phylogeny, our microbial isolate was designated *P. megaterium* SI1-IITJ for subsequent analyses. The Roary tool matrix indicated similarity of SI1-IITJ with other strains of *P. megaterium*. Comparative genomic analysis indicated that only a small percentage of genes were commonly shared among the studied strains while a major part of the genomes is made up of the shell and cloud (Fig. 2B). Further, the results obtained from the Proksee tool provided a detailed graphical representation of the genome organization of *P. megaterium* SI1-IITJ, with clearly marked features such as coding sequences (CDSs), tRNAs, rRNAs, and other functional elements. The map also included tracks representing the distribution of GC content and GC skew across the genome, offering insights into the genomic stability and nucleotide composition (Fig. 2C). Genes contributing to F^-^ stress tolerance, including F^-^ transporters and genes for plant growth promotion, viz., nitrogen fixation, auxin biosynthesis, phosphate solubilization, ACC deaminase activity, and siderophore biosynthesis, were identified from the genome of *P. megaterium* SI1-IITJ (Table 1). The genome analysis also revealed that SI1-IITJ contained genes for resistance to β-lactam and glycopeptide antibiotics (Table S2). However, the antibiotic resistance genes were not found in the mobilome of this strain, confirming its safety in biofertilizer applications (Table S3).

**Fig. 2.**
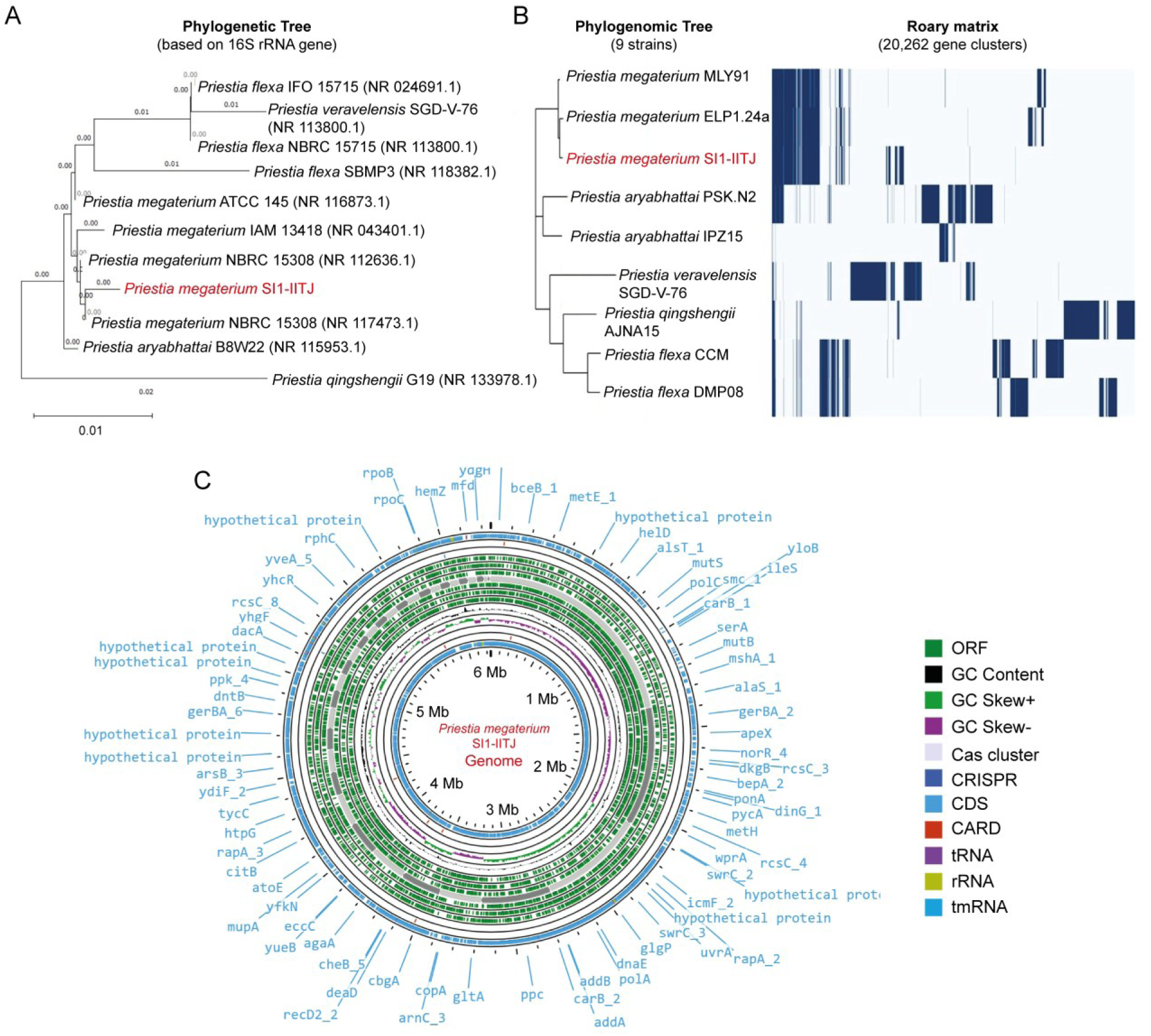
Phylogeny and genome analysis of *Priestia megaterium* SI1-IITJ. **(A)** A phylogenetic tree of *Priestia* species is shown. The tree was constructed from 16S rRNA gene sequences of the microbial isolate SI1-IITJ and its homologs, identified by BLASTN against the NCBI database, by the neighbor-joining method using MEGA-X. Values shown at the branch positions are the observed evolutionary distances. The *P. quigshengii* G19 16S rRNA gene sequence represented the out-group in the phylogenetic tree. **(B)** A phylogenomic tree of SI1-IITJ, constructed with the FastTree v2.1.10 tool via the Roary platform. The left panel represents the phylogenetic relationship of *P. megaterium* SI1-IITJ with other *Priestia* species. The right panel depicts the core and accessory genes shared by different *Priestia* species. **(C)** Genome map of *P. megaterium* SI1-IITJ. The circular genome map of the microbial isolate was generated with the Proksee online tool (https://proksee.ca/) using the complete genome sequence of the isolate and annotated by the PROKKA program.

**Table 1.**
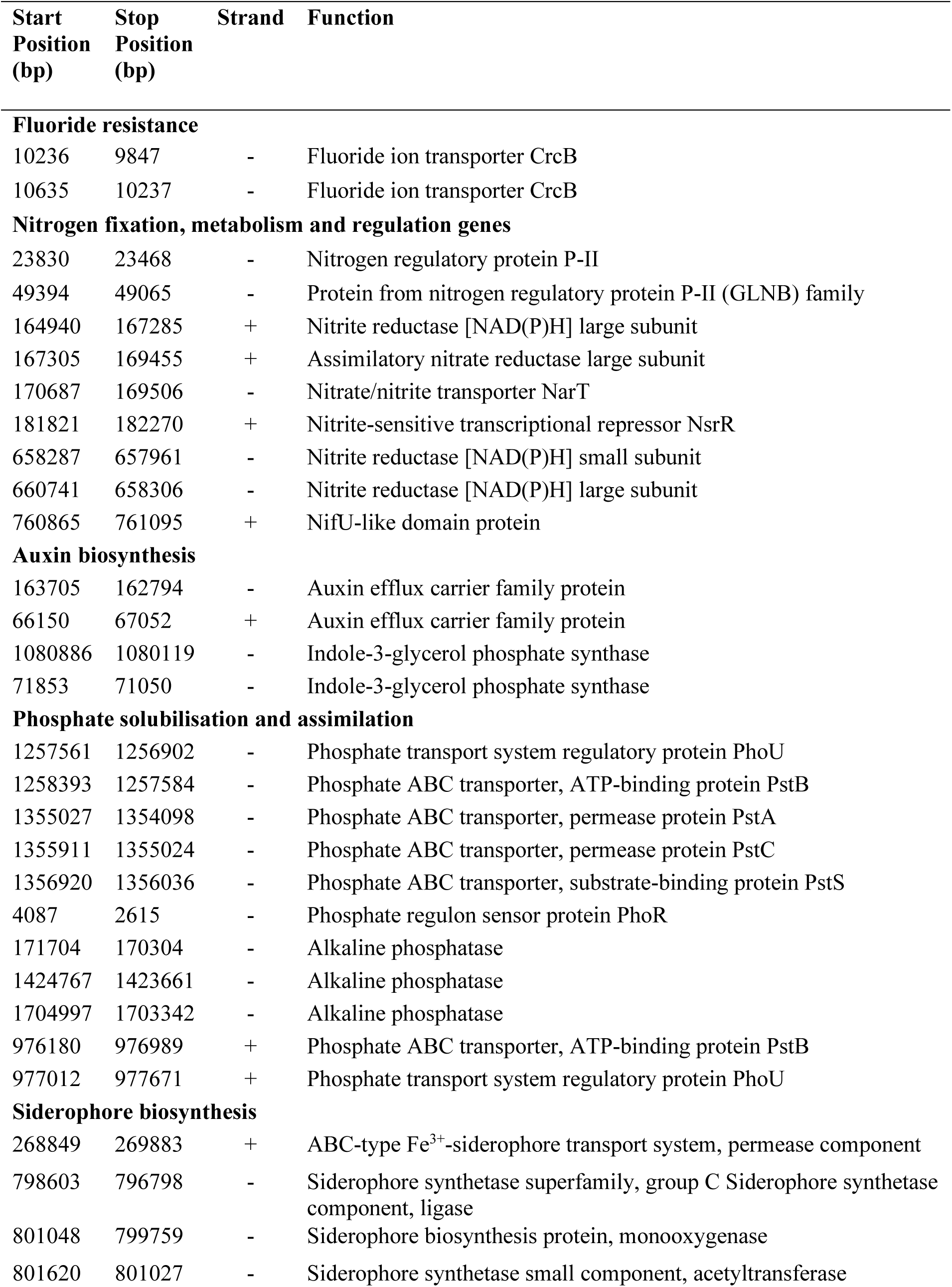
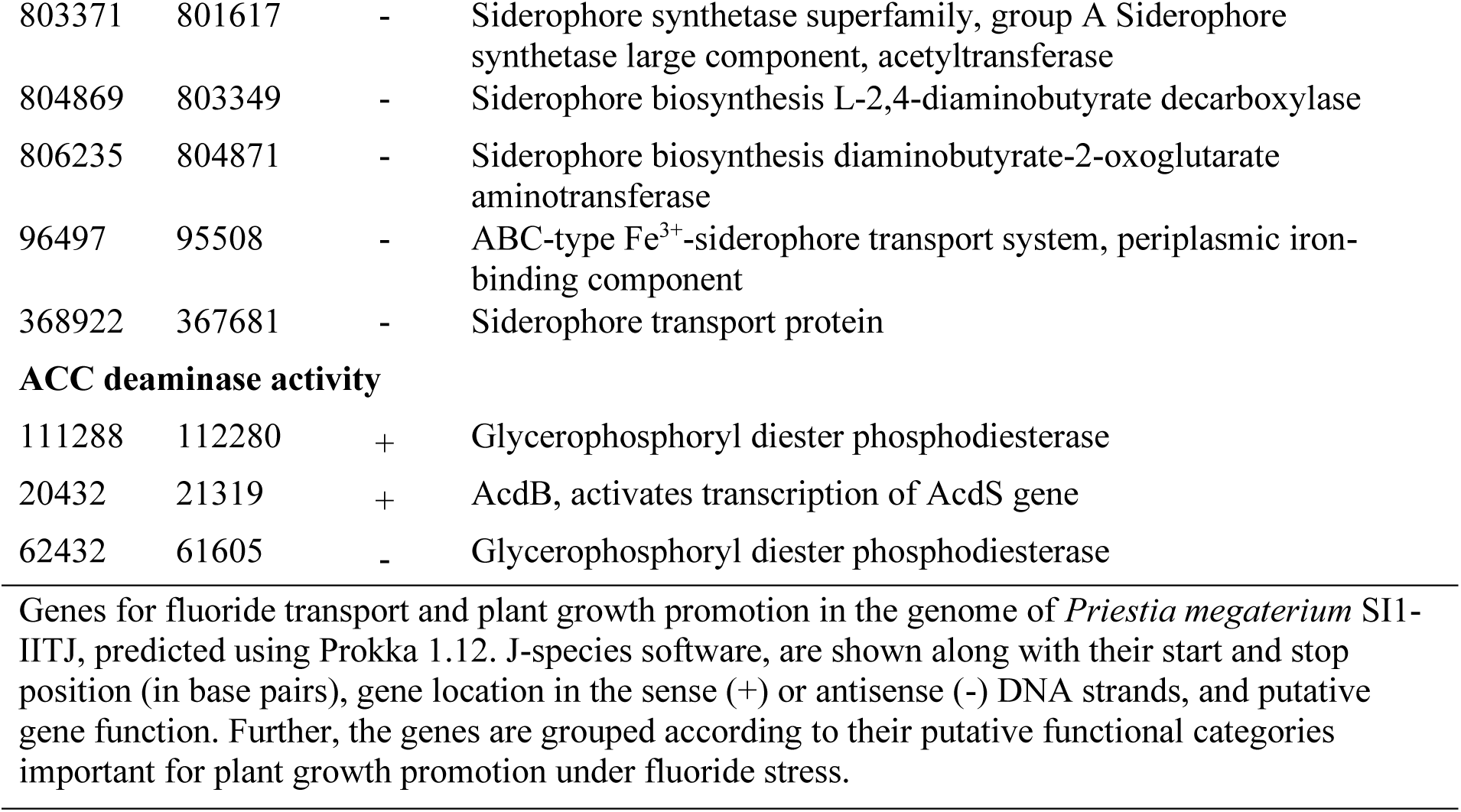
*Priestia megaterium* SI1-IITJ genes for fluoride tolerance and plant growth promotion.

| **Start Position (bp)** | **Stop Position (bp)** | **Strand** | **Function** |
| --- | --- | --- | --- |
| **Fluoride resistance** |  |  |  |
| 10236 | 9847 | - | Fluoride ion transporter CrcB |
| 10635 | 10237 | - | Fluoride ion transporter CrcB |
| **Nitrogen fixation, metabolism and regulation genes** |  |  |  |
| 23830 | 23468 | - | Nitrogen regulatory protein P-II |
| 49394 | 49065 | - | Protein from nitrogen regulatory protein P-II (GLNB) family |
| 164940 | 167285 | + | Nitrite reductase [NAD(P)H] large subunit |
| 167305 | 169455 | + | Assimilatory nitrate reductase large subunit |
| 170687 | 169506 | - | Nitrate/nitrite transporter NarT |
| 181821 | 182270 | + | Nitrite-sensitive transcriptional repressor NsrR |
| 658287 | 657961 | - | Nitrite reductase [NAD(P)H] small subunit |
| 660741 | 658306 | - | Nitrite reductase [NAD(P)H] large subunit |
| 760865 | 761095 | + | NifU-like domain protein |
| **Auxin biosynthesis** |  |  |  |
| 163705 | 162794 | - | Auxin efflux carrier family protein |
| 66150 | 67052 | + | Auxin efflux carrier family protein |
| 1080886 | 1080119 | - | Indole-3-glycerol phosphate synthase |
| 71853 | 71050 | - | Indole-3-glycerol phosphate synthase |
| **Phosphate solubilisation and assimilation** |  |  |  |
| 1257561 | 1256902 | - | Phosphate transport system regulatory protein PhoU |
| 1258393 | 1257584 | - | Phosphate ABC transporter, ATP-binding protein PstB |
| 1355027 | 1354098 | - | Phosphate ABC transporter, permease protein PstA |
| 1355911 | 1355024 | - | Phosphate ABC transporter, permease protein PstC |
| 1356920 | 1356036 | - | Phosphate ABC transporter, substrate-binding protein PstS |
| 4087 | 2615 | - | Phosphate regulon sensor protein PhoR |
| 171704 | 170304 | - | Alkaline phosphatase |
| 1424767 | 1423661 | - | Alkaline phosphatase |
| 1704997 | 1703342 | - | Alkaline phosphatase |
| 976180 | 976989 | + | Phosphate ABC transporter, ATP-binding protein PstB |
| 977012 | 977671 | + | Phosphate transport system regulatory protein PhoU |
| **Siderophore biosynthesis** |  |  |  |
| 268849 | 269883 | + | ABC-type Fe<sup>3+</sup>-siderophore transport system, permease component |
| 798603 | 796798 | - | Siderophore synthetase superfamily, group C Siderophore synthetase component, ligase |
| 801048 | 799759 | - | Siderophore biosynthesis protein, monooxygenase |
| 801620 | 801027 | - | Siderophore synthetase small component, acetyltransferase |
| 803371 | 801617 | - | Siderophore synthetase superfamily, group A Siderophore synthetase large component, acetyltransferase |
| 804869 | 803349 | - | Siderophore biosynthesis L-2,4-diaminobutyrate decarboxylase |
| 806235 | 804871 | - | Siderophore biosynthesis diaminobutyrate-2-oxoglutarate aminotransferase |
| 96497 | 95508 | - | ABC-type Fe<sup>3+</sup>-siderophore transport system, periplasmic iron-binding component |
| 368922 | 367681 | - | Siderophore transport protein |
| **ACC deaminase activity** |  |  |  |
| 111288 | 112280 | + | Glycerophosphoryl diester phosphodiesterase |
| 20432 | 21319 | + | AcdB, activates transcription of AcdS gene |
| 62432 | 61605 | - | Glycerophosphoryl diester phosphodiesterase |

Genes for fluoride transport and plant growth promotion in the genome of *Priestia megaterium* SI1-IITJ, predicted using Prokka 1.12. J-species software, are shown along with their start and stop position (in base pairs), gene location in the sense (+) or antisense (-) DNA strands, and putative gene function. Further, the genes are grouped according to their putative functional categories important for plant growth promotion under fluoride stress.

### *Priestia megaterium* SI1-IITJ enhances plant development under fluoride stress and non-stressed environments

Inoculation of *P. megaterium* SI1-IITJ to *A. thaliana* seedlings growing in hydroponics led to a significant improvement of the root lengths in two weeks under both control (0 mM) and 1 mM NaF stress (Fig. 3A). Similar improvements were observed in a F^-^-sensitive ecotype Star-8 (68.2 ± 12.9% under 0 mM and 70.6 ± 12.6% under 1 mM NaF) and a F^-^-tolerant ecotype Col-0 (50.8 ± 4.7% under 0 mM and 27.7 ± 2.6% under 1 mM NaF) of *A. thaliana* (Fig. 3B). In Star-8, the rosette diameters also increased significantly (42.9 ± 3.8% under 0 mM and 70.5 ± 8.6% under 1 mM NaF) due to SI1-IITJ (Fig. 3B). During a 30-day soil culture, irrigation with higher concentrations (50‒150 mM) of NaF led to a visible increase of rosette size in pots inoculated with SI1-IITJ (Fig. 3C). The same observation was reflected in a significant 70.5 ± 0.5% increase in shoot biomass and a 51.9 ± 23% increase in root biomass under 100 mM NaF stress (Fig. 3D). Similar observations were obtained for both Col-0 (Fig. 3C, D) and Star-8 (Fig. S2). These results indicate that SI1-IITJ can increase the growth of *A. thaliana* under F^-^ stress.

**Fig. 3.**
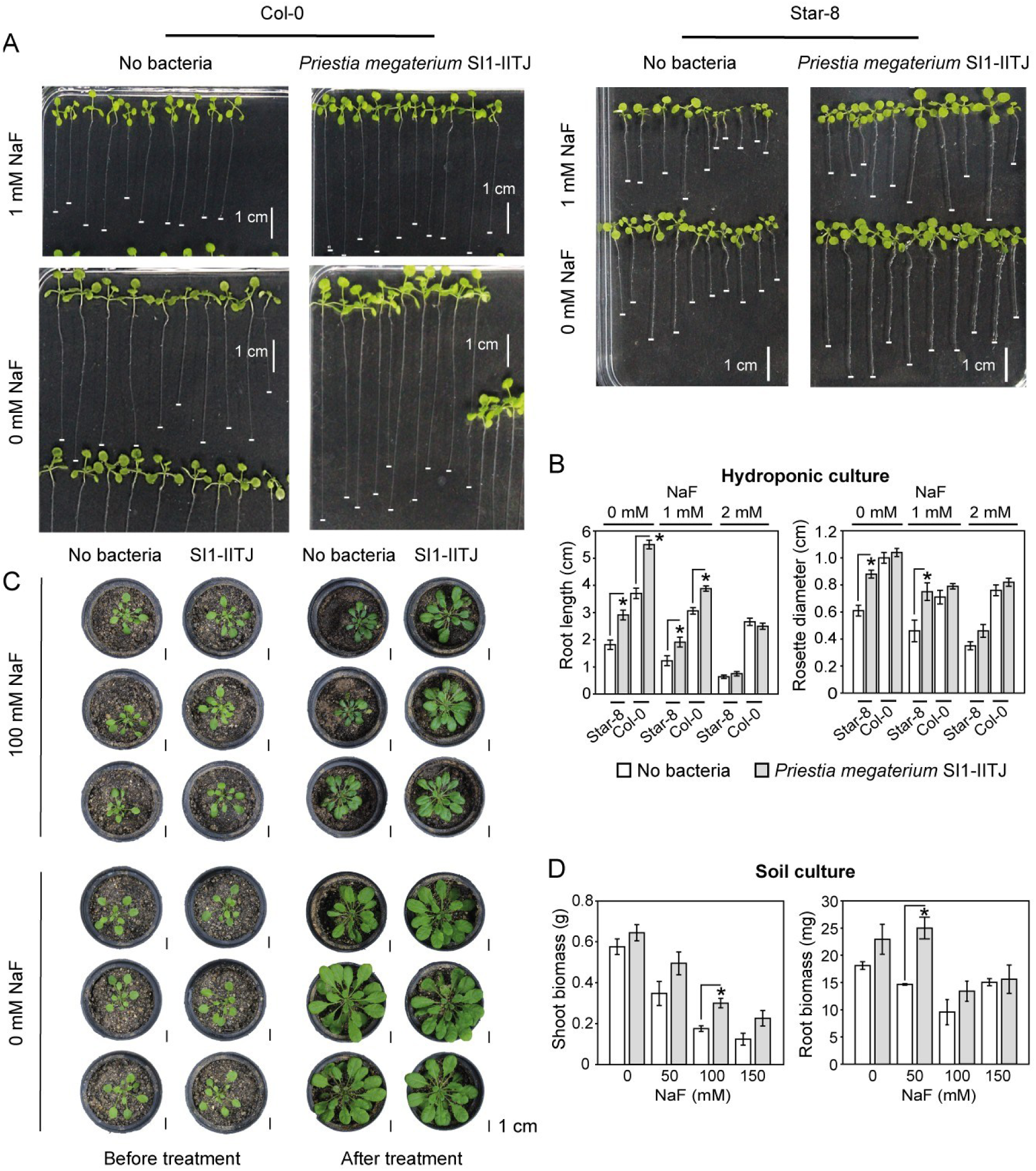
Plant growth promotion of *Priestia megaterium* SI1-IITJ under fluoride stress. **(A)** Growth of *Arabidopsis thaliana* ecotype Col-0 and Star-8 seedlings grown hydroponically for two weeks in ¼-Hoagland’s solution with or without 1 mM NaF, in the presence or absence of *P. megaterium* SI1-IITJ, is shown. The hydroponic media were inoculated with sterile PBS (no bacteria) or *P. megaterium* SI1-IITJ (see Methods). The seedlings were placed on solidified agar and photographed from an equal distance. The root tip positions are marked by white horizontal bars. The white vertical bars represent 1 cm. **(B)** The average root lengths and rosette diameters are shown with standard error after two weeks of hydroponic growth under 0, 1, and 2 mM NaF. **(C)** The effects of *P. megaterium* SI1-IITJ on one-month-old Col-0 plants in the soil culture experiment are shown. The pots were irrigated with different concentrations of NaF (see Methods). Three biological replicates are shown for each treatment. Black bars represent a scale of 1 cm **(D)** Average shoot and root biomass of ten plants in each treatment are presented with standard error after one-month growth in soil under 0, 50, 100, and 150 mM NaF. Asterisks indicate significant differences at *P* < 0.05 (Student’s *t*-test, *N* = 10).

### *Priestia megaterium* SI1-IITJ perturbs the global gene expression of *Arabidopsis thaliana* under fluoride stress

A transcriptome analysis of *A. thaliana* seedlings co-cultured with *P. megaterium* SI1-IITJ for two weeks under F^-^ stress indicated significant alterations in gene expression profiles, with 55 genes significantly induced (log₂fold-change > 1, *FDR* < 0.05) and 103 genes down-regulated (log₂FC < –1, *FDR* < 0.05) in SI1-IITJ-inoculated plants compared to non-inoculated plants (Fig. 4, Table 2).

**Fig. 4.**
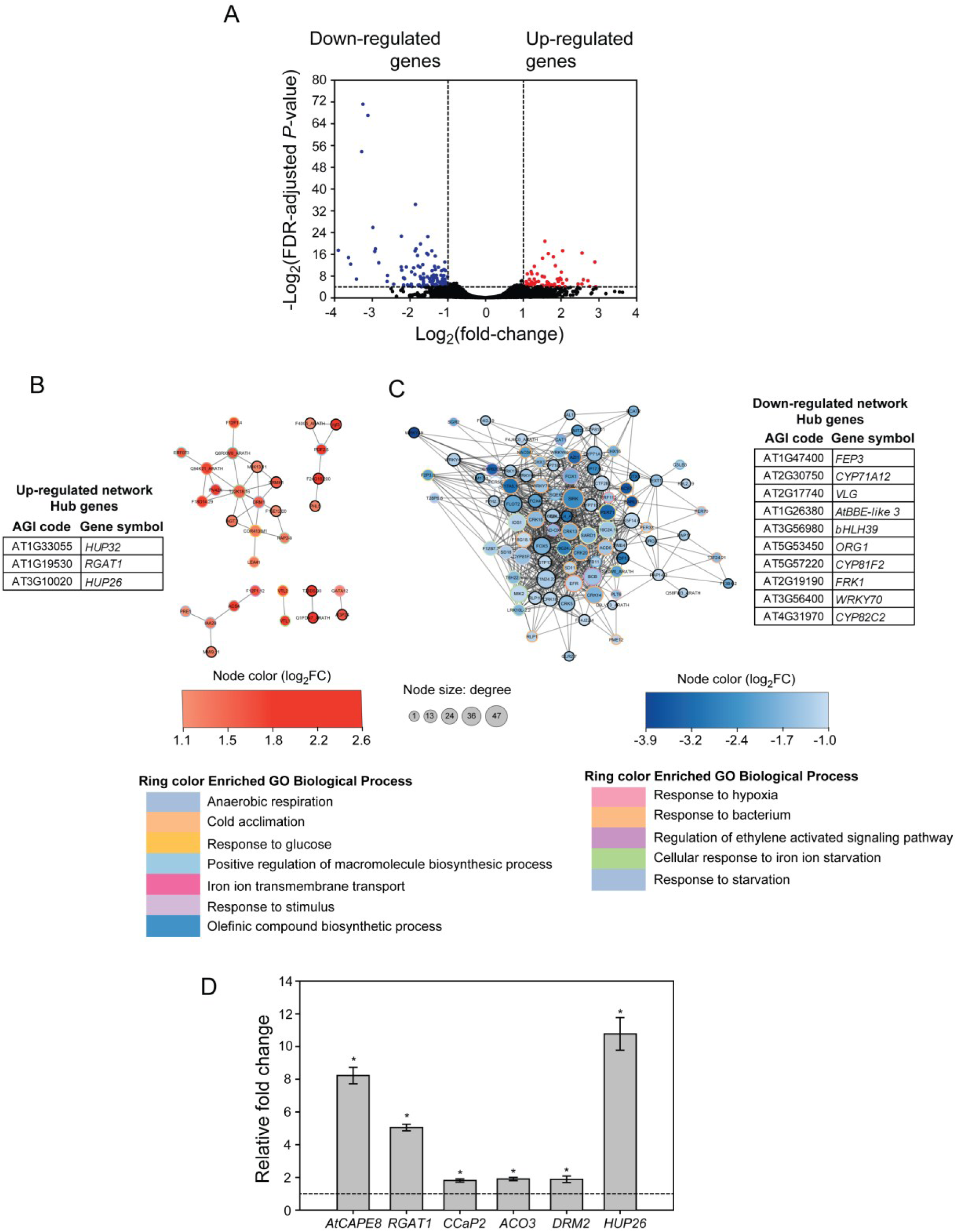
Transcriptome analysis of *Arabidopsis thaliana* after coculture with *Priestia megaterium* SI1-IITJ under fluoride stress. *Arabidopsis thaliana* seedlings were grown hydroponically for two weeks in ¼-Hoagland’s solution with 1 mM NaF without or with *P. megaterium* SI1-IITJ. **(A)** A volcano plot of SI1-IITJ-inoculated versus non-inoculated samples is shown. The Y-axis represents -log_2_(adjusted *P*-value), or Benjamini-Hochberg *false discovery rate* (*FDR*)-adjusted *P*-value (Fisher’s exact test, *N* = 3 biological replicates), and the X-axis represents -log_2_(fold-change) of expression levels between SI1-IITJ-inoculated and non-inoculated samples. The horizontal dotted lines delimit an *FDR* < 0.05, whereas the vertical dotted lines delineate −1 > -log_2_(FC) > 1. Up-(log_2_FC >1, *FDR* < 0.05) and down-regulated (log_2_FC <-1, *FDR* < 0.05) genes are denoted by red and blue dots, respectively. Protein-protein interaction networks of upregulated genes (**B**) and down-regulated genes (**C**), identified by STRING and visualized through Cytoscape. Red nodes represent the up-regulated genes, whereas the down-regulated genes are represented by blue nodes, with color shades denoting the log_2_(FC). The hub genes in the networks are shown along with their Arabidopsis Genome Initiative (AGI) codes and gene symbols. The node size key representing the degree of the nodes is shown below. The colored rings around the nodes represent the enriched Gene Ontology (GO) biological process categories, with color keys depicted below. (**D**) Real-time qPCR validation of six randomly selected genes, viz., *AtCAPE8*, *RGAT1*, *CCaP2*, *ACO3*, *DRM2*, and *HUP26*, up-regulated in the transcriptome analysis, is shown. Each gene’s expression levels were compared between SI1-IITJ-inoculated and non-inoculated samples. The expression levels were further normalized using *AtUBQ1* as the internal standard. The dotted line shows the level of gene expression in non-inoculated samples. Asterisks indicate significant changes in expression levels between SI1-IITJ-inoculated and non-inoculated samples (Student’s *t*-test, *P* < 0.05; *N* = 3).

**Table 2.**
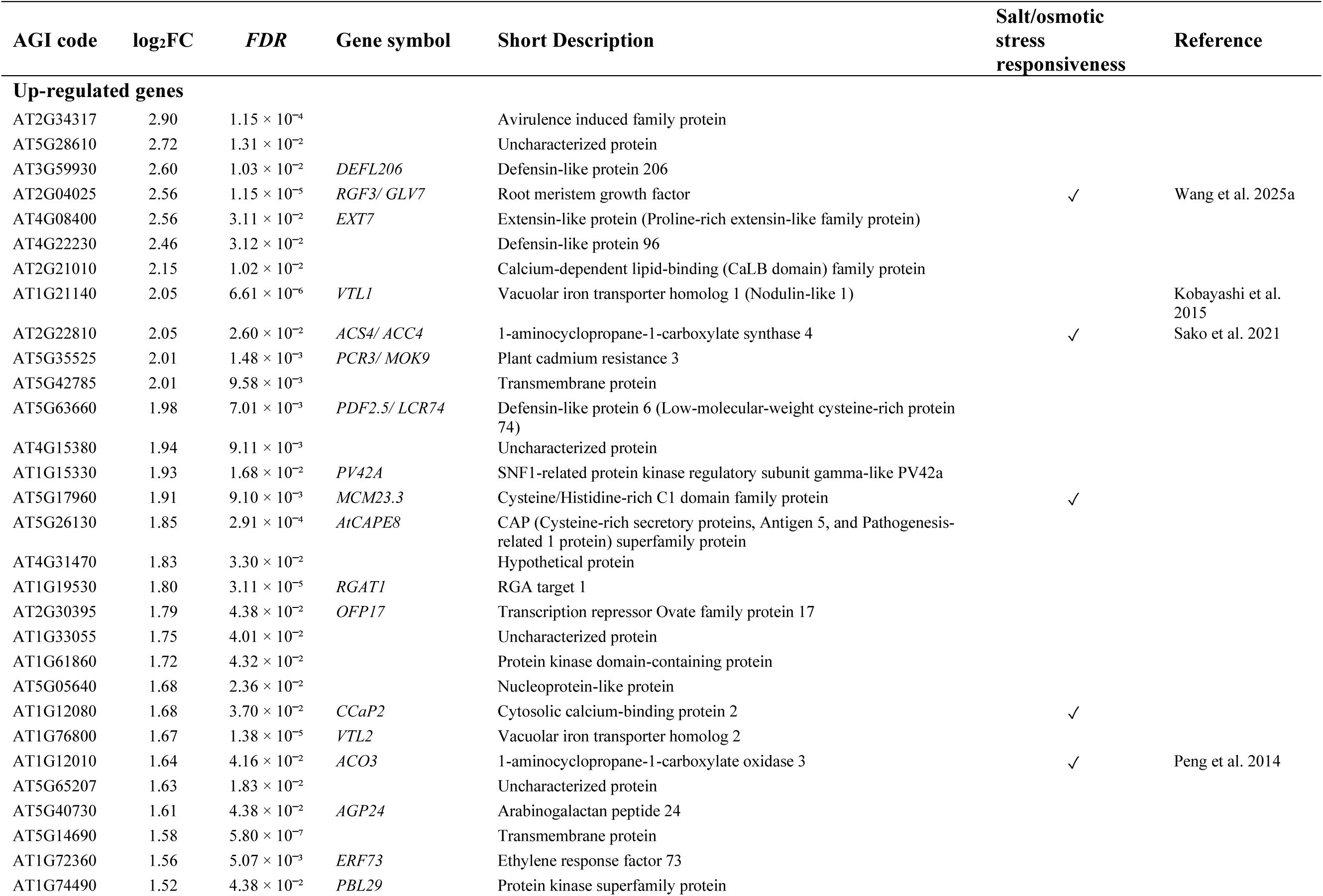
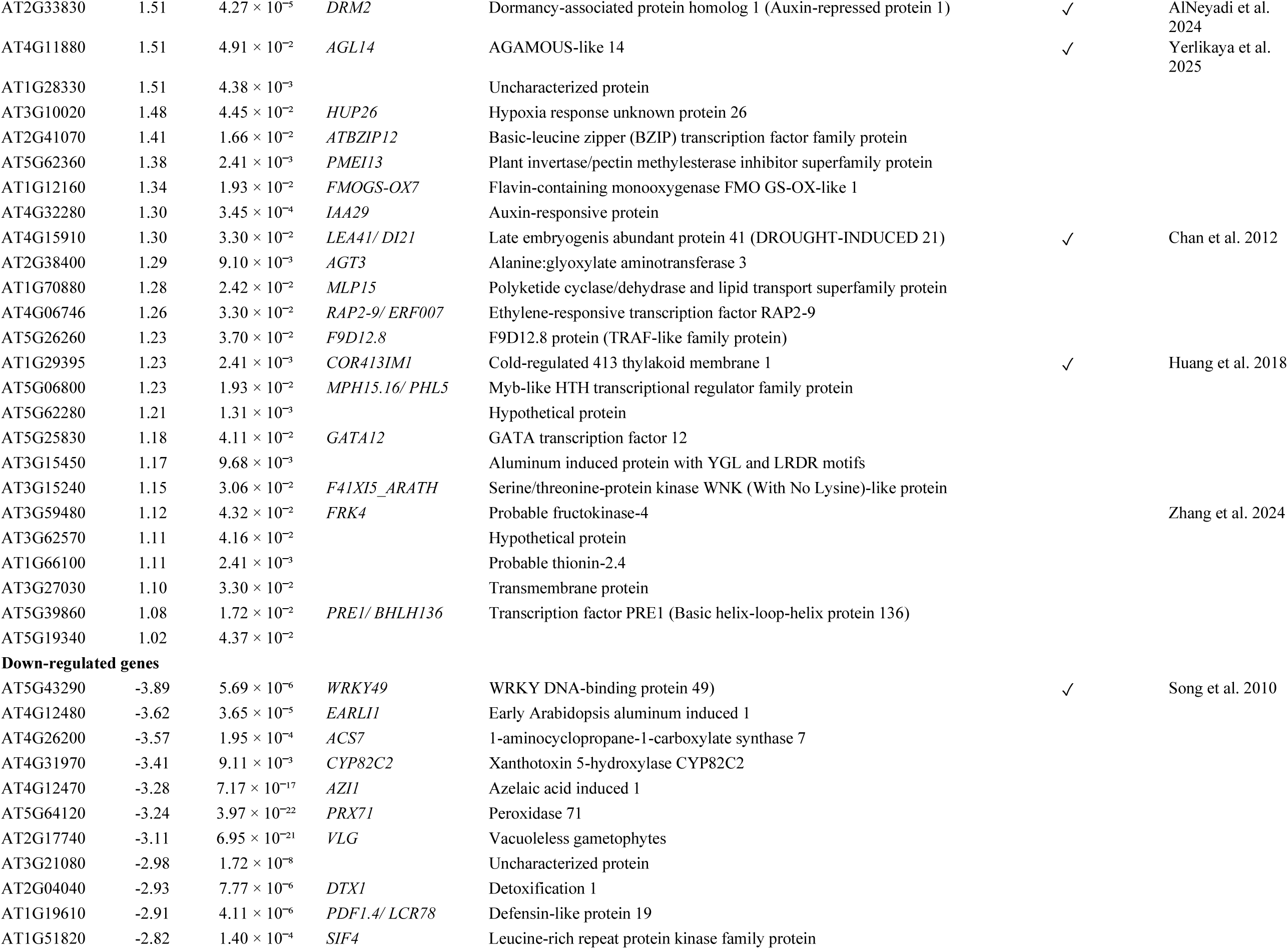
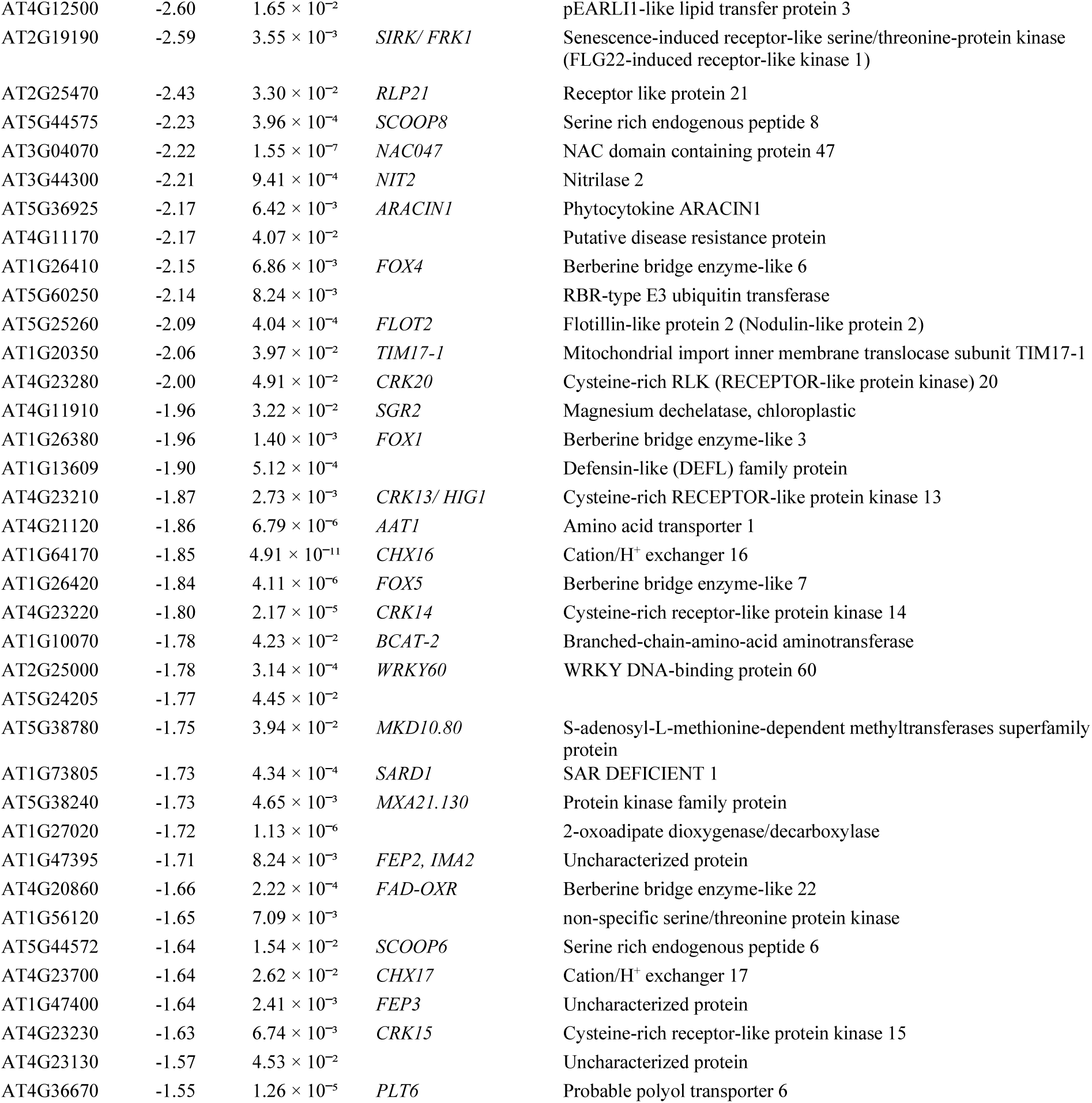
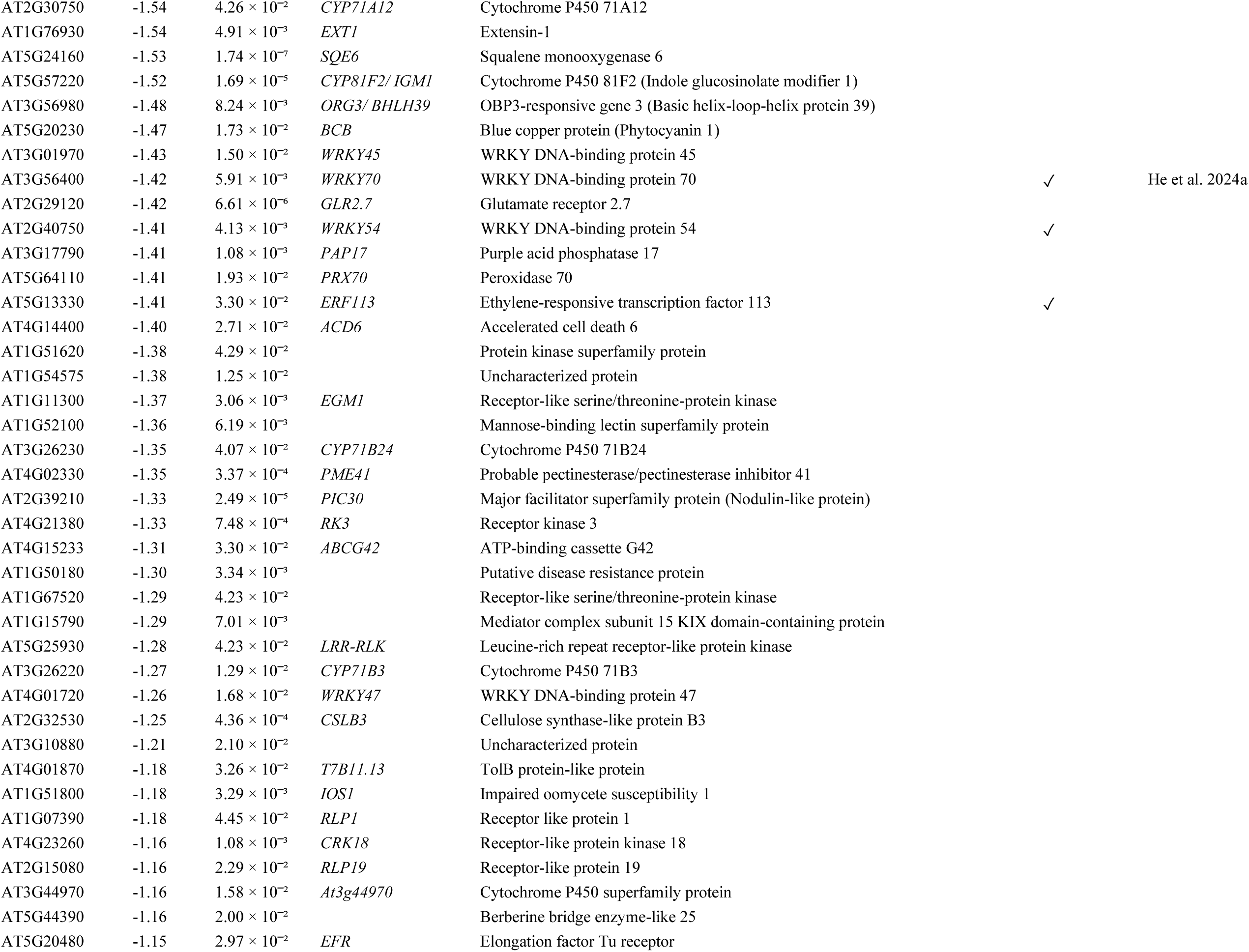
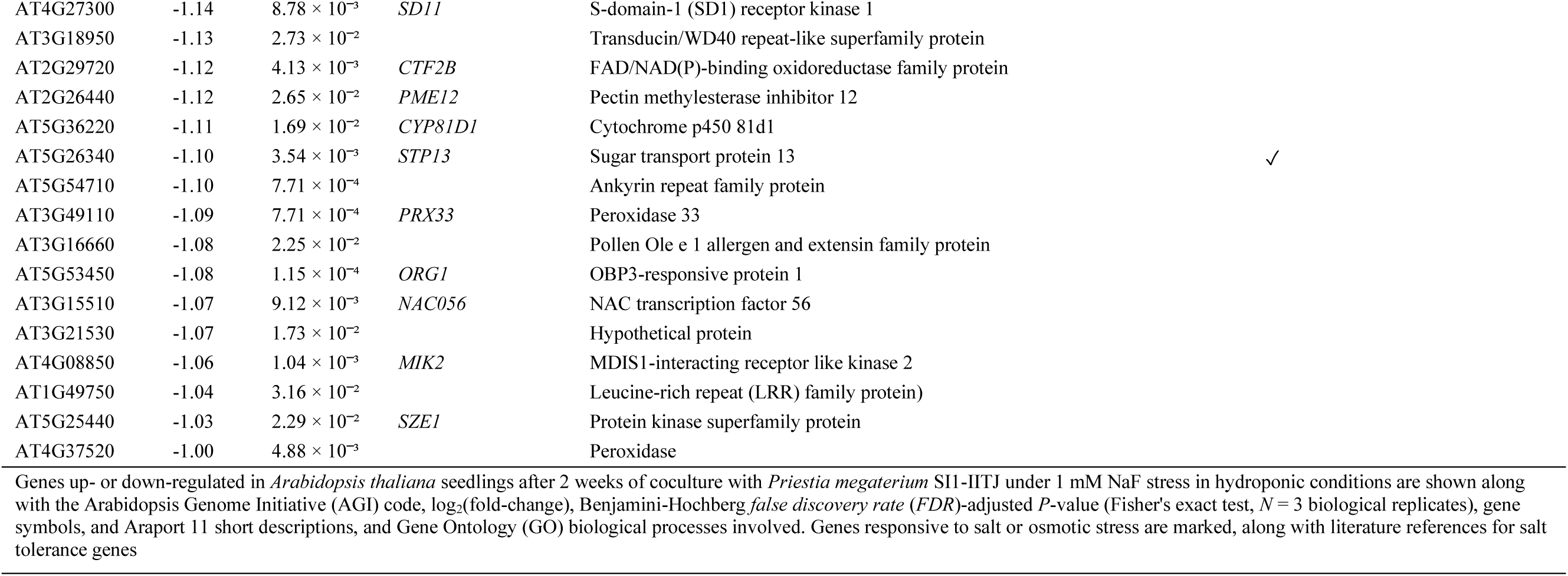
*Arabidopsis thaliana* genes differentially expressed after coculture with *Priestia megaterium* SI1-IITJ under fluoride stress.

Functional enrichment analysis of 53 up-regulated genes revealed their involvement in the biological processes, viz., response to stimulus, response to abiotic stimulus, iron (Fe) and manganese (Mn) transmembrane transport, intracellular Fe sequestration, olefinic compound biosynthesis, regulation of root meristem growth, and anaerobic respiration. An enriched Kyoto Encyclopedia of Genes and Genomes (KEGG) pathway included ‘cysteine and methionine metabolism’ (Table 3). These up-regulated genes belong to critical physiological and developmental processes in plants, regulating growth and stress adaptation. Genes associated with cell wall development and modification showed induction due to *P. megaterium* SI1-IITJ, including *extensin 7* (*EXT7*; AT4G08400), *pectin methyl-esterase inhibitor 13* (*PMEI13*; AT5G62360), and *arabinogalactan protein 24* (*AGP24*; AT5G40730). A significant number of 13 up-regulated genes involved in responses to abiotic stress, including high salt (NaCl) and water deprivation, were significantly induced, including *plant cadmium resistance 3* (*PCR3*; AT5G35525), *cysteine/histidine-rich c1 domain family protein* (*MCM23.3*; AT5G17960), *cytosolic calcium-binding protein 2* (*CCaP2*; AT1G12080), *drought-induced 21* (*DI21*; AT4G15910), *cold regulated 314 inner membrane 1* (*COR413IM1*; AT1G29395), and *dormancy/auxin associated family protein 2* (*DRM2*; AT2G33830). Genes associated with anaerobic respiration and hypoxia response, *hypoxia response unknown protein 26* (*HUP26*; AT3G10020) and *HUP32* (AT1G33055), were also induced. Many genes were also associated with phytohormones: ethylene and auxin biosynthesis and signaling pathways, including *1-amino-cyclopropane-1-carboxylic acid (ACC) oxidase 3* (*ACO3*; AT1G12010), *ACC synthase* (*ACS4*; AT2G22810), *ethylene response factor 73* (*ERF73*; AT1G72360), *ERF007* (AT4G06746), *indole-3-acetic acid inducible 29* (*IAA29*; AT4G32280), and *paclobutrazol resistance 1* (*PRE1*; AT5G39860). Several defensin-like proteins, such as *DEFL206* (AT3G59930), *cysteine-rich receptor-like linase 15* (*CRK15*; AT4G22230), *plant defensin 2.5* (*PDF2.5*; AT5G63660), and *plant thionin* (*F15E12.20*; AT1G66100), were also induced. Transcription factors (TFs) regulating root meristem growth, *root growth factor 3* (*RGF3*; AT2G04025) and *agamous-like 14* (*AGL14*; AT4G11880), chloroplast development-regulating MYB-like helix-turn-helix TF *phosphate response1-like 1* (*PHL5*; AT5G06800), and *GATA transcription factor 12* (*GATA12*; AT5G25830), involved in regulating chlorophyll biosynthesis, were among the induced genes. In addition, Fe transporter genes responsible for maintaining Fe and Mn homeostasis were prominently represented among the induced genes, including *vacuolar iron transporter 1* (*VTL1*; AT1G21140) and *VTL2* (AT1G76800), underscoring their role in nutrient regulation under F^-^ stress. Up-regulated genes involved in metabolic pathways included *flavin-containing monooxygenase* (*FMOGS-OX7*; AT1G12160), involved in glucosinolate biosynthesis, and *fructokinase 4* (*FRK4*; AT3G59480), involved in fructose metabolism and starch biosynthesis.

**Table 3.**
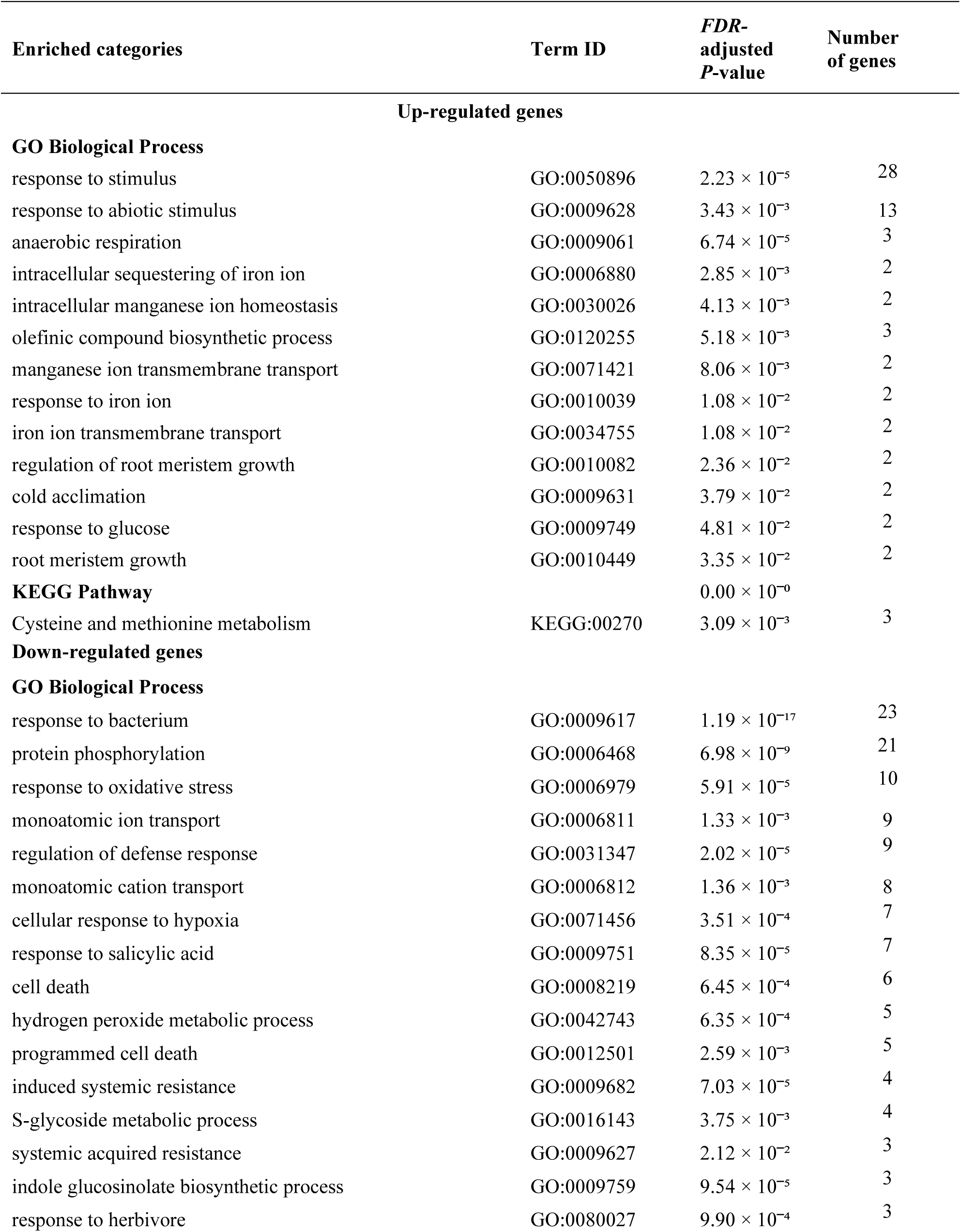
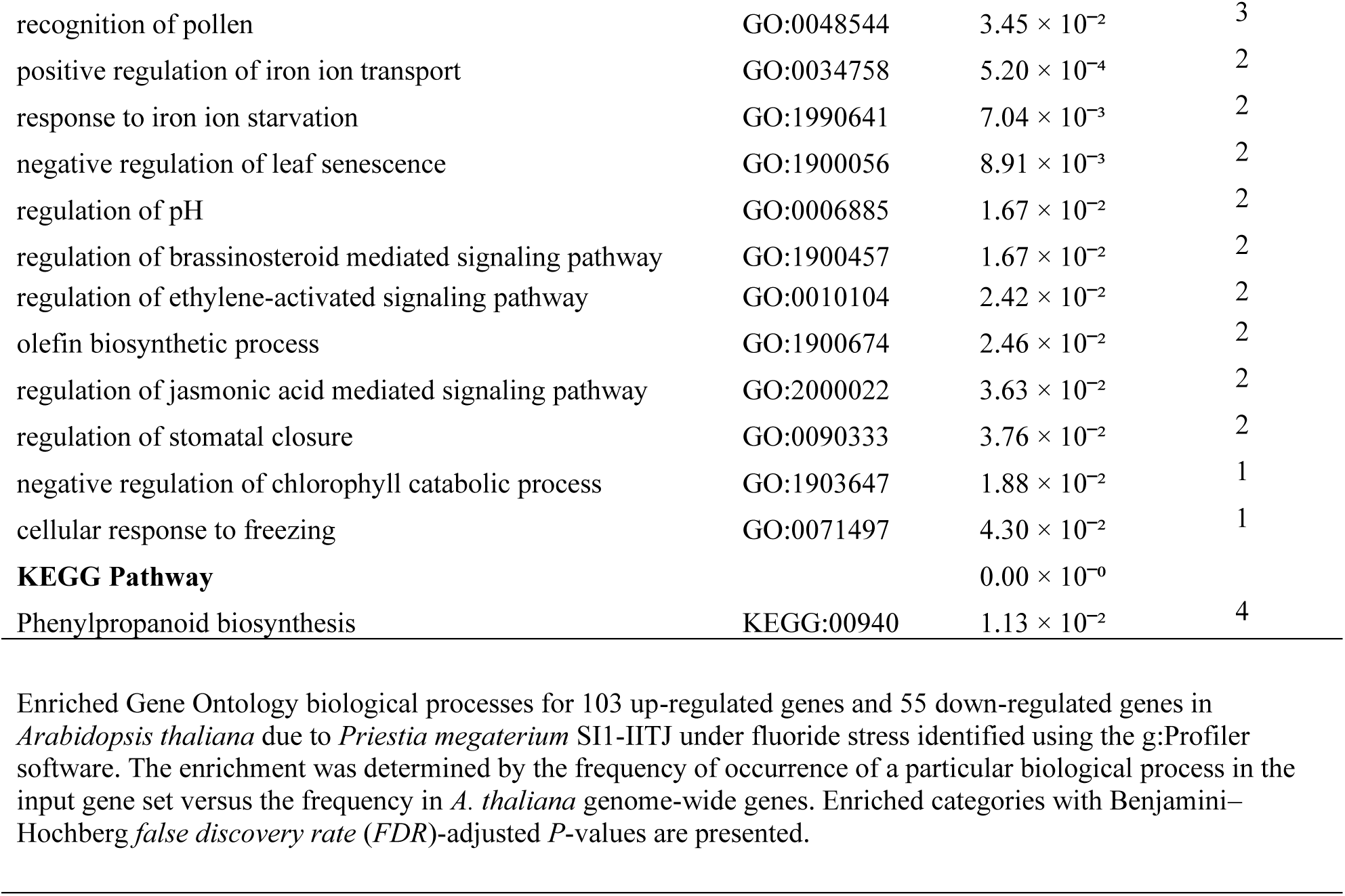
Enriched functional categories of genes differentially expressed in *Arabidopsis thaliana* upon coculture with *Priestia megaterium* SI1-IITJ under fluoride stress.

| Enriched categories | Term ID | FDR-adjusted<br>P-value | Number<br>of genes |
| --- | --- | --- | --- |
| **Up-regulated genes** |  |  |  |
| **GO Biological Process** |  |  |  |
| response to stimulus | GO:0050896 | $2.23 \times 10^{-5}$ | 28 |
| response to abiotic stimulus | GO:0009628 | $3.43 \times 10^{-3}$ | 13 |
| anaerobic respiration | GO:0009061 | $6.74 \times 10^{-5}$ | 3 |
| intracellular sequestering of iron ion | GO:0006880 | $2.85 \times 10^{-3}$ | 2 |
| intracellular manganese ion homeostasis | GO:0030026 | $4.13 \times 10^{-3}$ | 2 |
| olefinic compound biosynthetic process | GO:0120255 | $5.18 \times 10^{-3}$ | 3 |
| manganese ion transmembrane transport | GO:0071421 | $8.06 \times 10^{-3}$ | 2 |
| response to iron ion | GO:0010039 | $1.08 \times 10^{-2}$ | 2 |
| iron ion transmembrane transport | GO:0034755 | $1.08 \times 10^{-2}$ | 2 |
| regulation of root meristem growth | GO:0010082 | $2.36 \times 10^{-2}$ | 2 |
| cold acclimation | GO:0009631 | $3.79 \times 10^{-2}$ | 2 |
| response to glucose | GO:0009749 | $4.81 \times 10^{-2}$ | 2 |
| root meristem growth | GO:0010449 | $3.35 \times 10^{-2}$ | 2 |
| **KEGG Pathway** | | $0.00 \times 10^{-0}$ | |
| Cysteine and methionine metabolism | KEGG:00270 | $3.09 \times 10^{-3}$ | 3 |
| **Down-regulated genes** |  |  |  |
| **GO Biological Process** |  |  |  |
| response to bacterium | GO:0009617 | $1.19 \times 10^{-17}$ | 23 |
| protein phosphorylation | GO:0006468 | $6.98 \times 10^{-9}$ | 21 |
| response to oxidative stress | GO:0006979 | $5.91 \times 10^{-5}$ | 10 |
| monoatomic ion transport | GO:0006811 | $1.33 \times 10^{-3}$ | 9 |
| regulation of defense response | GO:0031347 | $2.02 \times 10^{-5}$ | 9 |
| monoatomic cation transport | GO:0006812 | $1.36 \times 10^{-3}$ | 8 |
| cellular response to hypoxia | GO:0071456 | $3.51 \times 10^{-4}$ | 7 |
| response to salicylic acid | GO:0009751 | $8.35 \times 10^{-5}$ | 7 |
| cell death | GO:0008219 | $6.45 \times 10^{-4}$ | 6 |
| hydrogen peroxide metabolic process | GO:0042743 | $6.35 \times 10^{-4}$ | 5 |
| programmed cell death | GO:0012501 | $2.59 \times 10^{-3}$ | 5 |
| induced systemic resistance | GO:0009682 | $7.03 \times 10^{-5}$ | 4 |
| S-glycoside metabolic process | GO:0016143 | $3.75 \times 10^{-3}$ | 4 |
| systemic acquired resistance | GO:0009627 | $2.12 \times 10^{-2}$ | 3 |
| indole glucosinolate biosynthetic process | GO:0009759 | $9.54 \times 10^{-5}$ | 3 |
| response to herbivore | GO:0080027 | $9.90 \times 10^{-4}$ | 3 |
| recognition of pollen | GO:0048544 | $3.45 \times 10^{-2}$ | 3 |
| positive regulation of iron ion transport | GO:0034758 | $5.20 \times 10^{-4}$ | 2 |
| response to iron ion starvation | GO:1990641 | $7.04 \times 10^{-3}$ | 2 |
| negative regulation of leaf senescence | GO:1900056 | $8.91 \times 10^{-3}$ | 2 |
| regulation of pH | GO:0006885 | $1.67 \times 10^{-2}$ | 2 |
| regulation of brassinosteroid mediated signaling pathway | GO:1900457 | $1.67 \times 10^{-2}$ | 2 |
| regulation of ethylene-activated signaling pathway | GO:0010104 | $2.42 \times 10^{-2}$ | 2 |
| olefin biosynthetic process | GO:1900674 | $2.46 \times 10^{-2}$ | 2 |
| regulation of jasmonic acid mediated signaling pathway | GO:2000022 | $3.63 \times 10^{-2}$ | 2 |
| regulation of stomatal closure | GO:0090333 | $3.76 \times 10^{-2}$ | 2 |
| negative regulation of chlorophyll catabolic process | GO:1903647 | $1.88 \times 10^{-2}$ | 1 |
| cellular response to freezing | GO:0071497 | $4.30 \times 10^{-2}$ | 1 |
| **KEGG Pathway** | | $0.00 \times 10^{-0}$ | |
| Phenylpropanoid biosynthesis | KEGG:00940 | $1.13 \times 10^{-2}$ | 4 |

Enriched Gene Ontology biological processes for 103 up-regulated genes and 55 down-regulated genes in *Arabidopsis thaliana* due to *Priestia megaterium* SI1-IITJ under fluoride stress identified using the g:Profiler software. The enrichment was determined by the frequency of occurrence of a particular biological process in the input gene set versus the frequency in *A. thaliana* genome-wide genes. Enriched categories with Benjamini–Hochberg *false discovery rate* (*FDR*)-adjusted *P*-values are presented.

Functional enrichment analysis of 103 down-regulated genes revealed their involvement in systemic acquired resistance (SAR), response to hypoxia, bacteria, oxidative stress, and Fe starvation, monoatomic ion transport, protein phosphorylation, indole glucosinolate biosynthesis, and regulation of brassinosteroid- and SA-mediated signaling pathways. Phenylpropanoid biosynthesis was the only KEGG pathway enriched among down-regulated genes (Table 3). Our results indicated broad suppression of transcriptional regulators integral to SAR and plant defense against bacteria, possibly to allow beneficial PGPR growth. For example, the stress-related TFs *WRKY45* (AT3G01970), *WRKY47* (AT4G01720), *WRKY49* (AT5G43290), *WRKY54* (AT2G40750), *WRKY60* (AT2G25000), and *WRKY70* (AT3G56400) were found to be down-regulated. Transporter genes *detoxification 1* (*DTX1*; AT2G04040), *cation/H^+^ exchanger 16* (*CHX16*; AT1G64170), *CHX17* (AT4G23700), *sugar transport protein 13* (*STP13*; AT5G26340), and *amino acid transporter 1* (*AAT1*; AT4G21120), involved in cellular detoxification, ionic balance, and nutrient allocation, showed reduced expression. Repression of *Fe-uptake-inducing peptide 2* (*FEP2*; AT1G47395) and *FEP3* (AT1G47400), both associated with Fe deficiency signaling, was observed. Genes linked to cell wall organization and remodeling, including *extensin-1* (*EXT1*; AT1G76930), *probable pectinesterase inhibitor 41* (*PME41*; AT4G02330), and *cellulose synthase-like protein B3* (*CSLB3*; AT2G32530), were down-regulated, potentially reflecting shifts in cell wall plasticity or defense fortification. Genes involved in secondary metabolism, such as *cytochrome P450 protein family 82, subfamily C2* (*CYP82C2*; AT4G31970), *CYP81F2* (AT5G57220), *squalene monooxygenase 6* (*SQE6*; AT5G24160), and others in the cytochrome P450 family, contributing to biotic resistance, were repressed. Several receptor-like kinases, e.g., *EF-Tu receptor* (*EFR*; AT5G20480), *FLG22-induced receptor-like kinase 1* (*FRK1*; AT2G19190), and members of the CRK family, involved in early pathogen recognition and downstream signaling pathways, were also down-regulated. Defense-associated genes, including *azelaic acid induced 1* (*AZI1*; AT4G12470), *early Arabidopsis aluminum induced 1* (*EARLI1*; AT4G12480), *systemic acquired resistance deficient 1* (*SARD1*; AT1G73805), *accelerated cell death 6* (*ACD6*; AT4G14400), *impaired oomycete susceptibility 1* (*IOS1*; AT1G51800), *plant defensin 1.4* (*PDF1.4*; AT1G19610), and *defensin-like (DEFL) family protein* (AT1G13609) were among the most significantly repressed, revealing the extensive downshifting of induced systemic resistance mechanisms. Multiple peroxidases involved in ROS metabolism and cell wall lignification during defense, such as *peroxidase 33* (*PRX33*; AT3G49110), *PRX71* (AT5G64120), *PRX70* (AT5G64110), and AT4G37520, were down-regulated. In contrast to *ACS4* up-regulation, *ACS7* (AT4G26200), another isoform of *ACC synthase*, was strongly down-regulated, as was the ethylene-responsive TF *ERF113* (AT5G13330). The auxin biosynthesis gene *nitrilase 2* (*NIT2*; AT3G44300) was also repressed.

A network analysis was conducted to understand the molecular interactions underlying the F^-^ resistance conferred by *P. megaterium* to *A. thaliana* seedlings (Fig. 4B, C). A comparative overview of the networks showed that the down-regulated PPI network had more DEGs, while the up-regulated network consisted of fewer genes. A functional analysis of the up-regulated network genes revealed enriched biological processes such as anaerobic respiration, cold acclimation, response to glucose, positive regulation of macromolecule biosynthetic processes, Fe ion transmembrane transport, response to stimulus, and olefinic compound biosynthetic processes (Fig. 4B). Among the up-regulated DEGs, two highly interconnected subnetworks were identified, each comprising five genes with more than one interaction. These subnetworks reflect crucial molecular modules linked to phytohormone signaling, stress responses, and developmental regulation under the influence of PGPRs. The first subnetwork included two ethylene signaling genes, *ACO3* and *ACS4*, auxin signaling gene *IAA29*, a bHLH TF *PRE1*, and *PMEI13*, a pectin methylesterase inhibitor involved in regulating root growth (Chen et al. 2018). The first sub-network of genes promoted stress-responsive growth by modulating the crucial phytohormones auxin and ethylene, and cell wall remodelling. The second subnetwork included an abiotic stress-responsive gene *with-no-lysine (WNK)-like protein* (*F41XI5_ARATH*; AT3G15240), root growth-regulating TF *RGF3*, a chloroplast development-regulating TF *PHL5*, and the defensin genes *PDF2.5* and *DEFL206*, also responsive to various abiotic stresses (Domingo et al. 2024). Altogether, up-regulated genes of the second sub-network are linked to developmental regulation, both defense and abiotic stress responses, and signaling pathways modulated by PGPRs. Key hub genes identified in the up-regulated network include defense, ROS-regulating, and hypoxia-responsive genes *HUP26* and *HUP32* (Huh et al. 2020), and a high salt-responsive GA-signaling gene *RGA target 1* (*RGAT1*; AT1G19530), which are closely associated with salt stress mitigation under PGPR influence (Yang et al. 2023; Zhou et al. 2024). In the presence of PGPRs, the up-regulation of *HUP26* and *HUP32* is part of a suite of adaptive responses that may help the plant cope with localized oxygen depletion or metabolic stress induced by bacterial activity (Cantabella et al. 2022). The induced genes in the transcriptome, *AtCAPE8*, *RGAT-1*, *CCaP2*, *ACO3*, *DRM2*, and *HUP26* showed a similar trend of up-regulation when validated by qPCR (Fig. 4D).

In the more interconnected down-regulated gene network, the most significantly enriched GO biological processes included response to bacteria and hypoxia, regulation of ethylene signaling, and response to Fe starvation (Fig. 4C). Further analysis revealed 41 most-densely-connected central genes of the network, from which 10 hub genes were shortlisted based on their centrality scores (Fig. 4C). The down-regulated hub genes included *FEP3*, a small peptide involved in Fe deficiency signaling (Lichtblau et al. 2022); three cytochrome P450 genes, *CYP71A12* (AT2G30750), *CYP81F2* (AT5G57220), *CYP82C2* (AT4G31970), and *berberine bridge enzyme-like 3* (*AtBBE-like 3*; AT1G26380), involved in the biosynthesis of indole glucosinolates for defense against pathogens; two TFs, *bHLH39* (AT3G56980), involved in Fe deficiency responses, and *WRKY70* (AT3G56400), an activator of salicylic acid (SA) signaling defense genes and repressor of JA signaling genes; flagellin-induced genes *vacuoleless gametophytes* (*VLG*; AT2G17740), and the receptor kinase *FRK1*. Suppression of these hub defense genes may facilitate PGPR colonization (Stringlis et al. 2017). Interestingly, many of these genes are also responsive to salt and osmotic stress (Table 2). Another down-regulated hub gene, *OBP3-responsive gene 1* (*ORG1*; AT5G53450), is a lipid-binding kinase involved in root halotropism and osmotic stress response (Deolu-Ajayi et al. 2019).

### *Priestia megaterium* SI1-IITJ reduces fluoride and ROS, enhancing chlorophyll and nitrogen content in plant tissues

We explored the physiological basis of the observed phenotypic gains of *A. thaliana* upon inoculation with *P. megaterium* SI1-IITJ in the soil culture assay during F^-^ treatment (100 mM NaF) or non-stressful conditions (0 mM NaF). Interestingly, SI1-IITJ reduced the F^-^ content of plant rosettes by 64 ± 1.7% (Fig. 5A). We observed a 24.9 ± 0.4% increase in total chlorophyll content under non-stressful conditions and a 1.6 ± 0.1% increase under F^-^ stress due to SI1-IITJ (Fig. 5B). The H_2_O_2_ levels in plant tissues increased under F^-^ stress. However, SI1-IITJ helped to reduce a significant 48.9 ± 1% in H_2_O_2_ levels in plant tissues under F^-^ (Fig. 5C). As a PGPR, SI1-IITJ increased the total tissue nitrogen and phosphate by 18.7 ± 4.1% and 8.2 ± 1.5%, respectively, under non-stressful conditions, possibly by enhancing nutrient uptake (Fig. 5D, 5E). The enhancement of tissue nitrogen content due to SI1-IITJ was maintained in F^-^ stress by 30.4 ± 1.5% (Fig. 5D). The results suggest that SI1-IITJ increases plant growth under F^-^ stress by inhibiting F^-^ uptake, reducing oxidative stress, and enhancing nutrient mining (Fig. 6).

**Fig. 5.**
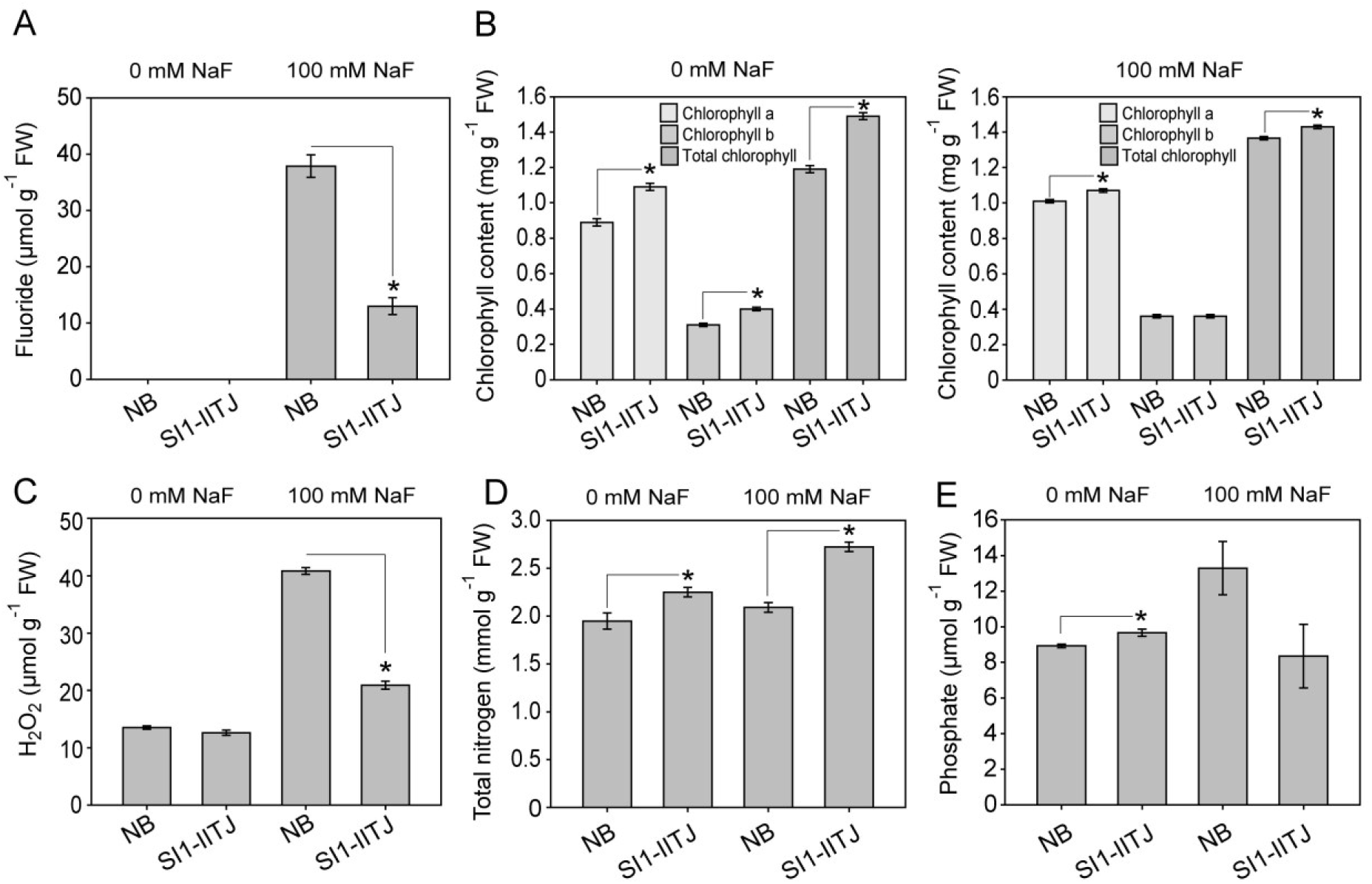
Plant fluoride, chlorophyll, nutrient, and ROS content after coculture with *Priestia megaterium* SI1-IITJ under fluoride stress. Rosette leaves of one-month-old *Arabidopsis thaliana* ecotype Col-0 plants inoculated with *P. megaterium* SI1-IITJ or no bacteria (NB) under non-stressful conditions (0 mM) or 100 mM NaF stress were harvested and subjected to different biochemical measurements as follows: **(A)** Fluoride estimated using a F^-^ ion sensitive probe **(B)** chlorophyll content **(C)** H_2_O_2_ content **(D)** total tissue nitrogen **(E)** tissue phosphate content (**B-E** measured spectrophotometrically; See Methods). All values are expressed per gram of rosette fresh weight (FW). Asterisks indicate significant differences between NB versus SI1-IITJ treatments of *A. thaliana* (*P* < 0.05, Student’s *t*-test, *N* = 3).

**Fig. 6.**
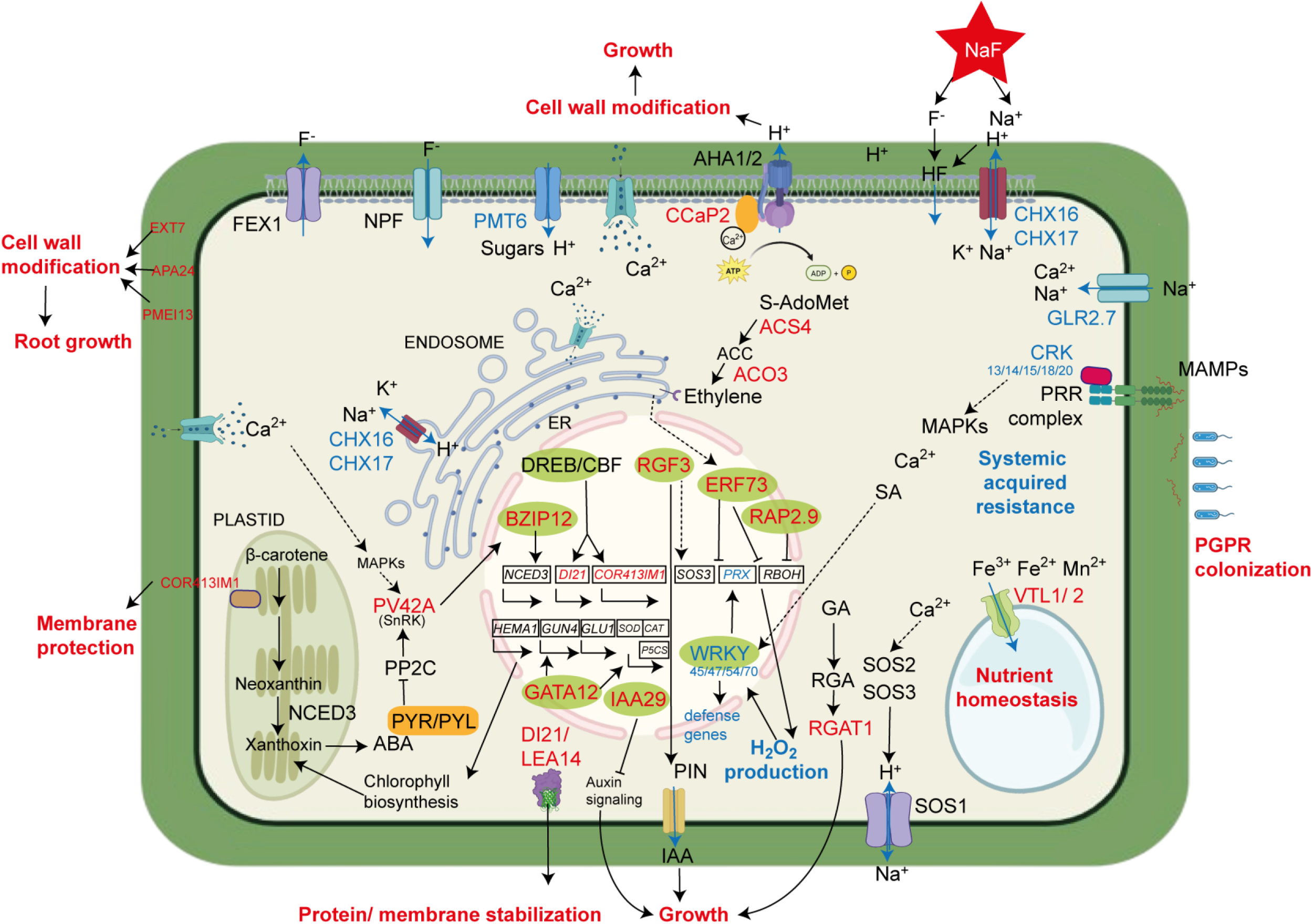
Mechanistic model of sodium fluoride stress mitigation in *Arabidopsis thaliana* by *Prestia megaterium* SI1-IITJ. Sodium fluoride (NaF) dissociates into Na*^+^* and F^-^ ions. F^-^ enters the cell through NPF transporters and combines with H^+^ to form lipophilic HF, entering passively through the cell membrane lipid bilayer. F^-^ is extruded from the cell by FEX1. Na^+^ can enter the cell through plasma membrane-localized cation/H^+^ antiporters CHX16 and CHX17 and the non-specific cation channel GLR2.7. Suppression of *CHXs* can reduce the H^+^ flow to the apoplast, reducing HF formation and its entry into the cell. CHX16/17, localizing at the endosomal membranes, regulates Na^+^/H^+^ homeostasis. Ca^2+^ entry in the cell from the apoplast or the endoplasmic reticulum (ER) under NaF stress leads to the activation of CCaP2, associating with the H^+^ ATPase, AHA1/2, leading to the efflux of H^+^, loosening the cell wall to promote growth. Ca^2+^ signals reach the SnRK, PV42A of the abscisic acid (ABA) signaling pathway, which regulates stress response transcription factors (TFs). The upregulated TF BZIP12 induces *NCED3* expression, leading to higher ABA production. GATA12 induces the expression of *HEMA1*, *GUN4*, and *GLU1* to enhance chlorophyll biosynthesis, and reactive oxygen species (ROS) detoxifying genes *CAT*, *SOD*, and *P5CS*. The protein/membrane-stabilizing chaperone DI21/LEA41 and the chloroplast membrane-protecting COR413IM1 are regulated by DREB/CBF TFs. RGF3 regulates the Ca^2+^ signaling gene SOS3, which regulates the Na^+^/H^+^ antiporter SOS1. RGF3 also regulates auxin efflux PIN proteins to regulate growth. IAA29, a negative regulator of auxin signa ling, further regulates growth. Ethylene is produced from S-adenosyl methionine (S-AdoMet) by the induced enzymes ACS4 and ACO3. Ethylene signaling reaches ERF73 in the nucleus through the ER. The ethylene-responsive TFs ERF73 and RAP2.9 negatively regulate RBOH and peroxidases (PRXs) to suppress H_2_O_2_ production. Microbe-associated molecular patterns (MAMPs) bind to pattern recognition receptor (PRR) complexes, incorporating CRK proteins, relaying the signal through MAPKs, Ca^2+^, and salicylic acid (SA) to WRKY TFs, which regulate defense genes of systemic acquired resistance (SAR). Suppression of *CRKs* and *WRKYs* leads to the suppression of *PRXs*, leading to reduced H_2_O_2_ production. Suppression of SAR leads to increased PGPR colonization. The induced gibberellic acid (GA) signaling gene *RGAT1* promotes growth. Vacuolar sequestration of Fe^2+^/Fe^3+^/Mn^2+^ by VTL1/2 leads to nutrient homeostasis and growth under NaF stress. Induction of the cell wall modifying enzymes, *PMEI13*, *EXT7*, and *APA24* leads to enhanced root growth. Black arrows indicate biochemical conversions or regulations, while blue arrows indicate transport. The genes and processes up-regulated by SI1-IITJ are shown in red, while the down-regulated ones are shown in blue. Check the main text for the full names and more details on the above genes. Figure created using BioRender (https://www.biorender.com/).

## Discussion

In the current study, we isolated *P. megaterium* SI1-IITJ from the root endospheres of Thar Desert plants and found that it could tolerate NaF up to 100 mM in LB media. Although F^-^ can passively enter the bacterial cell through the pH gradient across the membrane by the formation of HF (Marquis et al. 2003), F^-^ expulsion in *P. megaterium* SI1-IITJ possibly occurs through a specific F^-^ transporter, FluC/ CrcB, identified in its genome (Table 1). These small membrane proteins of the Fluc family are responsible for F^-^ tolerance in other bacteria, including *E. coli* and *Pseudomonas putida* (Last et al. 2016; Calero et al. 2022). Our F^-^ measurements indicated that at the stationary phase of bacterial growth (Fig. 1 E), most of the F^-^ was present outside the cell, whereas only a small amount was detected inside the bacteria (Fig. S1). This observation supported the expulsion of F^-^ outside the cell as the tolerance mechanism of SI1-IITJ. This is in contrast with another F^-^-tolerant PGPR, *Serratia sp.*, which could remove F^-^ from the culture medium, sequestering it inside the cell (Katiyar et al. 2024).

Subsequent studies with SI1-IITJ established its role as a PGPR, which promoted growth and NaF tolerance of *A. thaliana*. Both genomic sequence analysis and biochemical experiments indicated its PGPR properties, including phosphate solubilization, ACC deaminase activity, IAA production, and nitrogen assimilation, as predicted by the genome analysis. All the above traits were observed in the F^-^-tolerant PGPR *Serratia sp.* (Katiyar et al. 2024). Other F^-^-tolerant bacteria also show nitrate-reduction capabilities (Chellaiah et al. 2021; Thirumala et al. 2022). Different salt-tolerant PGPRs induce IAA production to promote root development, solubilize phosphate for improved nutrient availability, express ACC deaminase to reduce ethylene under stress, and help with nitrogen assimilation under high Na^+^ (Sharma et al. 2021; Xie et al. 2024). Although we detected siderophore biosynthesis genes in the genome of SI1-IITJ (Table 1), siderophore production in this strain could not be detected by biochemical tests, indicating either a nonfunctional gene cluster or environment-specific activity that we failed to mimic under laboratory conditions (Marik et al. 2024).

Although the role of PGPRs, viz. *Serratia sp.* and *Pseudomonas aeruginosa* in alleviating F^-^ stress in plants is well known (Katiyar et al. 2024, 2025), the precise molecular mechanisms of F^-^ toxicity alleviation in plants, particularly at the genetic level by PGPRs, are unknown, prompting us to conduct a comparative transcriptome analysis. Plants growing in the presence of SI1-IITJ showed distinct gene expression profiles compared to plants growing alone under NaF. We investigated the function of the perturbed genes to unravel the mode of action of SI1-IITJ. We employed the soluble sodium salt of F^-^, viz., NaF, to infuse the stress media in all our experiments, like earlier studies (Thirumala et al. 2022; Singh et al. 2024; Katiyar et al. 2024, 2025). In natural environments as well, higher concentrations of F^-^ are associated with high Na^+^ concentrations (Rafique et al. 2009). Particularly in the Thar Desert, the source of SI1-IITJ, high Na^+^ concentrations in soil are reported (Mehta et al. 1970). Hence, the plants were subjected to a combination of high Na^+^ and F*^-^*. F^-^ levels were significantly lowered in plant tissues by SI1-IITJ (Fig. 5A). Root-to-shoot transport of F^-^, as well as F^-^ efflux from the plant cell, occurs through a highly specific membrane-localized F^-^ exporter, FEX1 (Tausta et al. 2024). But we failed to detect any expression level changes of *AtFEX1* (AT2G41705) in our transcriptome, prompting us to analyze the expression patterns of possible non-specific F^-^ transporter genes. A plasma membrane-localized nitrate/peptide transporter, CsNPF2.3, from tea plant roots was found to transport F^-^ non-specifically (Niu et al. 2025). The *A. thaliana* orthologs of *NPF* are known to confer salt tolerance by translocating nitrate from root to shoot (Taochy et al. 2015). Although *NPF* genes were unchanged in our transcriptome results, we observed an enriched functional category, ‘monoatomic ion transport’, among the down-regulated genes, indicating a suppression of transporter genes in the transcriptome (Table 3). Among these genes, *GLR2.7* (AT2G29120) acts as a non-selective cation channel allowing entry of Ca^2+^ and Na^+^ into the cytoplasm (Davenport 2002), and *PMT6* (AT4G36670) helps in H^+^-symport of various substances, including sugars. Suppression of these genes (Table 2) by SI1-IITJ may indicate a possible attempt to prevent the entry of Na^+^ inside the cell. On the other hand, the suppression of *CHX16* and *CHX17*, Na^+^/H^+^ and K^+^/H^+^ antiporters involved in endosomal sequestration of Na^+^ ions and maintaining Na^+^/K^+^ homeostasis under salt stress (Chanroj et al. 2011), could be an indirect result of lowering the Na^+^ concentration in the cell under NaF by SI1-IITJ. CHX17 is also found to localize in the plasma membrane, having roles in apoplastic pH regulation (Sze et al. 2004; Chanroj et al. 2013). Hence, suppression of *CHX17* may disrupt H^+^ outflow to the apoplast, leading to less protonation of F^-^ ions, preventing HF formation and its entry into the cytoplasm.

Many of the DEGs in response to SI1-IITJ were responsive to high salt and osmotic stresses (Table 1). Notably, other strains of *P. megaterium* improved plant tolerance to high salt and drought stresses (Vilchez et al. 2018; Shalaby 2023; Zhu et al. 2023; Thakur et al. 2024). Among the genes up-regulated due to SI1-IITJ, the fructose metabolism gene *FRK4* was found in a salt tolerance quantitative trait locus (QTL) in *Brassica napus* (Zhang et al. 2024). The Fe and Mn transporter *VTL1* was identified in a genome-wide association study (GWAS) for salt stress tolerance. In that study, *VTL1* expression level variation as well as amino acid polymorphisms were significantly associated with high salt tolerance (Kobayashi et al. 2016). *VTL1* might help to alleviate high Na^+^ toxicity by regulating the Fe and Mn nutrient uptake and homeostasis in the cell (Gollhofer et al. 2014), apart from supplying nutrients to the symbiont PGPR (Walton et al. 2020). A homolog of *RGF3*, of the second up-regulated sub-network in our transcriptome (Fig. 4B), enhances salt tolerance by up-regulating the salt overly sensitive 3 (SOS3) pathway of salt tolerance in *Brassica napus* (Wang et al. 2025a). RGF3 plays a critical role in maintaining the root stem cell niche, a process heavily influenced by microbial interactions. RGF3 modulates auxin efflux proteins, regulating auxin distribution and root growth under stress (He et al. 2024b). Ectopic expression of a potato ortholog of *DRM2*, an auxin signaling gene induced by SI1-IITJ, leads to salt tolerance in *A. thaliana* (AlNeyadi et al. 2024). *DRM2* was also induced by a natural rhizobacterium of *A. thaliana*, *Pseudomonas* sp., promoting growth (Schwachtje et al. 2011). *PRE1*, a bHLH TF highly expressed in the roots in response to salt and drought stress, when overexpressed, enhanced root growth, promoting salt and drought tolerance (Du et al. 2023). *AGL14*, a MADS-box TF, up-regulated under salt stress in *A. thaliana*, also plays a role in root development and auxin signaling (Yerlikaya et al. 2025). PGPRs enhance root developmental programs by influencing hormonal signaling and activating root-specific transcriptional responses (Al-Turki et al. 2023). We observed induction of *PMEI13,* encoding a pectin methylesterase inhibitor involved in regulating root growth under salt stress through cell wall restructuring (Chen et al. 2018). Hence, induction of genes regulating auxin signaling root development under salt stress can very well explain the observed growth promotion of *A. thaliana* under NaF stress (Fig. 3). The *DI21* gene induced in our study is responsive to salt and is a late embryogenesis abundant (LEA) chaperone, protecting proteins from misfolding under salt stress (Chan et al. 2011). The chloroplast membrane-localized *COR413IM1* is a salt-responsive gene protecting chloroplast membranes from ROS-induced damage under salt stress. This gene is suppressed in the *salt hypersensitive mutant 9* (*sahy9*) mutant and regulated by CBF TFs (Huang et al. 2018). *CCaP2*, a plasma membrane-associated Ca²⁺-binding protein, decodes Ca²⁺ signals for stress signal transduction and interacts with H^+^-ATPases in the membrane, leading to apoplastic acidification loosening the cell wall to promote growth under stress (Kreps et al. 2002; Wang et al. 2024). *ERF73*, of the AP2/ERF transcription factor superfamily, is activated by diverse abiotic stresses such as drought, salinity, chilling, and heat, and modulates plant tolerance to various abiotic stresses (Ahmadizadeh et al. 2020; Wang et al. 2025b). *ATBZIP12* (AT2G41070) directly regulates the expression of *9-cis-epoxycarotenoid dioxygenase 3* (*NCED3*), a rate-limiting enzyme in abscisic acid (ABA) biosynthesis, and recent studies have shown that overexpression of *NCED3* enhances salt and drought stress tolerance in *A. thaliana*, suggesting that *ATBZIP12* may play a significant role in F^-^ stress adaptation by modulating ABA biosynthesis in our study (Baek et al. 2020; Truong et al. 2021; Bader et al. 2023). ABA, on the other hand, induces the expression of *Phaseolus vulgaris 42-kilodalton protein A* (*PV42A*; AT1G15330), an SnRK1 kinase, activating the transcription of salt-responsive genes (Wang et al. 2023). *PV42A* was also induced in the presence of SI1-IITJ, suggesting the involvement of ABA signaling to protect against salinity stress induced by NaF (Table 2). Overexpression of *GATA12*, induced by SI1-IITJ, enhanced photosynthesis, transpiration, and stomatal conductance under salinity and osmotic stress (Zhu et al. 2024). *MCM23.3* encodes a hormone-responsive signaling protein that is critical in mediating plant responses to salt stress by regulating gene expression, osmotic adjustment, and ion homeostasis (Bhaskar et al. 2015). The GA-signaling gene *RGAT1*, encoding the catalytic subunit A of DNA polymerase epsilon, regulates drought stress tolerance (Refaiy et al. 2025). On the other hand, down-regulation of some negative regulators of salt tolerance in our transcriptome can add to the NaF tolerance of *A. thaliana* in the presence of SI1-IITJ (Table 2). These included *WRKY49* and *WRKY70*, whose suppression increased salt tolerance (Song et al. 2010; He et al. 2024a).

Ethylene signaling modulates salinity stress responses through Na^+^/K^+^ balance, microtubule reassembly, and ROS homeostasis. Upon triggering ethylene production, salinity tolerance was enhanced in *A. thaliana* (Sako et al. 2021). The strong induction of ethylene biosynthesis genes *ACO3* and *ACS4* of the first up-regulated subnetwork can provide plant tolerance under NaF stress. Another PGPR, *Enterobacter* sp. SA187 induces salt tolerance in *A. thaliana* by activating the plant’s ethylene signaling pathway through ethylene synthesis via ACC oxidase enzymes such as ACO3 (Peng et al. 2014). When ethylene signaling or biosynthesis was disrupted, plants lost the PGPR-conferred salt tolerance, confirming the essential role of ethylene and ACO enzymes in this process (de Zélicourt et al. 2018). Again, mutants of petunia with edited *ACO* genes were salt sensitive (Naing et al. 2022). The ethylene signaling genes belonged to the enriched GO biological process ‘olefinic compound biosynthetic process,’ and the enriched KEGG pathway ‘cysteine and methionine metabolism.’ Olefinic compounds protect photosynthetic tissues from oxidative damage induced by various abiotic stresses, including high salt (Yeshi et al. 2022). Whereas ethylene signaling regulates olefinic compound biosynthesis starting from cysteine and methionine precursors (Bajguz et al. 2023).

The chlorophyll content of *A. thaliana* increased in the presence of SI1-IITJ under F^-^ stress (Fig. 5B), like other F^-^-tolerant PGPRs, *Pseudomonas aeruginosa* and *Serratia* sp. (Katiyar et al. 2024; Singh et al. 2024). This indicates that SI1-IITJ induces plant gene expression related to chlorophyll biosynthesis, also known for other PGPRs (Hanifah et al. 2023). In our transcriptome, chlorophyll biosynthesis-regulating gene *GATA12* (Bi et al. 2005) was induced, and *SGR2*, involved in chlorophyll degradation (Yamatani et al. 2022), was down-regulated (Table 3), which can explain a net increase in the observed chlorophyll content in SI1-IITJ-inoculated plants (Fig. 5B). A higher chlorophyll content may contribute to maintaining photosynthesis levels under stress, leading to enhanced growth of SI1-IITJ-inoculated plants (Katiyar et al. 2024; Marik et al. 2024).

SI1-IITJ inoculation led to significantly higher tissue nutrient levels, viz., phosphate and nitrogen content in *A. thaliana* under control conditions. The higher N levels were also maintained under F^-^ stress (Fig. 5D, E). This is possibly due to more efficient nutrient uptake caused by SI1-IITJ. Other salt-tolerant PGPR bacteria are known to increase phosphate and nitrogen in plants (Xie et al. 2024). Concomitant with our observed increase in tissue nitrogen content of *A. thaliana* by SI1-IITJ (Fig. 5D, E), we observed suppression of genes responsive to nitrogen starvation (Table 2). This included *GLR2.7*, a glutamate receptor, and *AAT1*, an amino acid transporter, both reported to be down-regulated under nitrogen treatment (Dey et al. 2025). A sugar transport protein, *STP13*, up-regulated under nitrogen starvation (Cun et al. 2024), was down-regulated in our study (Fig. 5E). An opposite pattern of down-regulation of nitrogen-starvation-responsive genes in our transcriptome indicated a condition of nitrogen excess in plants cocultured with SI1-IITJ (Dey et al. 2025). Similarly, we noticed an up-regulation of *VTL* genes, possibly helping Fe and Mn uptake and sequestration, and a down-regulation of the hub genes *FEP2* and *FEP3,* small peptides involved in Fe deficiency signaling (Lichtblau et al. 2022). PGPRs like *Paraburkholderia phytofirmans* and *Azospirillum brasilense* can indirectly influence Fe availability and signaling, often reducing plant Fe levels due to competitive uptake in the rhizosphere (Orellana et al. 2022).

We observed an enrichment of the ‘SAR’ biological process among down-regulated genes, indicating that the PGPR SI1-IITJ actively suppresses the host immune system. SAR is an immune response initiated on recognition of pathogens, involving a complex signaling cascade (Vlot et al. 2020). A signature of SAR activation is the transcriptional control of defense genes via WRKY TFs binding to W-box elements on downstream gene promoters. WRKY54 and WRKY70 positively regulate SA-mediated defenses (Bakshi and Oelmüller 2014; Chen et al. 2021). Our results indicate coordinated repression of many central defense marker genes, including *WRKY54* and *WRKY70*; *resistance methylated gene 1* (*RMG1*; AT4G11170), a nucleotide-binding leucine-rich repeat (NB-LRR) disease resistance gene, and *PRX70*, genes that are induced by PAMPs and are direct targets of WRKY proteins (Halter et al. 2021). In addition, the defense response has an inherent connection with cellular metabolic status, and biotic stress results in the synthesis of ROS that act as secondary messengers (Kissoudis et al. 2014). Our results indicate that genes involved in this network of ROS and SA signaling were also suppressed by SI1-IITJ. These comprise *Arabidopsis racine-like antimicrobial peptide 1* (*ARACIN1*; AT5G36925), a H_2_O_2_-inducible protein (Neukermans et al. 2015), and *senescence-associated gene 14* (*SAG14*; AT5G20230), playing a role at the crossroads of ROS and SA signaling (Cui et al. 2020). Down-regulation of *CYP71A12* and *CYP81F2* by SI1-IITJ would impair the biosynthesis of indolic glucosinolates, potentially reducing the plant’s resistance to microbes (van de Mortel et al. 2012; Koprivova et al. 2019). This suggests that such down-regulation might be a plant response that facilitates PGPR colonization. Although initially activated via MAMP recognition, beneficial microbes such as the PGPR *Pseudomonas simiae* WCS417 actively suppress *FRK1* and related genes to avoid eliciting a strong immune response (Stringlis et al. 2017). Down-regulation of all these genes, of WRKY-mediated transcription, PAMP sensing, and ROS and SA signaling, collectively yields strong molecular evidence that SI1-IITJ treatment actively suppresses the plant’s SAR response and, in turn, reduces ROS levels, possibly protecting the tissues and enabling growth under NaF stress. In addition, an induction of *DRM2* (Table 2, Fig. 4), a negative regulator of SAR, can reinforce suppression of plant defense (Roy et al. 2020). This modulation is critical for establishing mutualistic interactions with the PGPR (Stringlis et al. 2017).

Our results show that levels of ROS, viz., H_2_O_2_, were significantly reduced in SI1-IITJ-inoculated plants under F^-^ stress (Fig. 5). This is in line with other F^-^-tolerant PGPRs, which mitigated F^-^ toxicity in rice and tomato through antioxidant defense (Singh et al. 2024; Katiyar et al. 2024, 2025). In our transcriptome results, ‘response to oxidative stress’ and ‘hydrogen peroxide metabolic process’ were among the enriched functional categories of down-regulated genes. Among these, genes encoding class III peroxidases involved in H_2_O_2_ production, viz., *PRX33* (AT3G49110), *PRX70*, *PRX71* (AT5G64120), and AT4G37520, are known to inhibit plant growth by inducing ROS accumulation (Welinder et al. 2002; Raggi et al. 2015; Kámán-Tóth et al. 2019). Other genes, *flavin adenine dinucleotide-linked oxidoreductase 1* (*FOX1*; AT1G26380), *FOX4* (AT1G26410), *FOX5* (AT1G26420), AT5G44390, and *berberine bridge enzyme 22* (*BBE22*; AT4G20860), also play an important role in producing H_2_O_2_ (Locci et al. 2019; Costantini et al. 2023). Induction of *ERF73* and *related to apetala 2.9* (*RAP2.9*; AT4G06746) by SI1-IITJ may also reduce H_2_O_2_ levels as it negatively regulates *respiratory burst oxidase homolog* (*RBOH*) and several ROS-producing peroxidases (Yang et al. 2011; Yang et al. 2017). Suppression of ROS-producing genes can explain our biochemical detection of lower H_2_O_2_ levels in SI1-IITJ-inoculated plants, contributing to plant growth promotion under F^-^ stress. On the other hand, an up-regulated TF *GATA12* in our transcriptome led to the transcriptional up-regulation of antioxidant enzymes such as *catalase* (*CAT*), *superoxide dismutase* (*SOD*), *peroxidase* (*PER*), and *Δ¹-pyrroline-5-carboxylate synthetase* (*P5CS*), thereby strengthening the ROS scavenging system, in a previous study (Zhu et al. 2024).

In summary, our study indicates that *P. megaterium* SI1-IITJ, a root endophyte of Thar Desert plants, mitigates NaF toxicity by the induction of plant growth-promoting and abiotic stress-responsive genes, many of which belong to known molecular pathways of high salt tolerance (Fig. 6). Non-specific Na^+^ transporters were down-regulated, leading to possible prevention of Na^+^ entry in the cell. Lowering of F^-^ content in plant tissues by SI1-IITJ could be the result of diluting out the F^-^ ions per unit tissue weight due to enhanced growth and resultant plant biomass, or the prevention of its entry through the cell membrane as HF, by apoplastic pH regulation. Other mechanisms of growth promotion by this bacterium include enhancing chlorophyll biosynthesis and nutrient mining while suppressing genes of SAR, plant defense, and H_2_O_2_ production. This is the first report providing mechanistic insights on the F^-^ stress mitigation by PGPRs through the analysis of global plant gene expression. Future genetic studies, including reverse genetic analysis through gene knockout and complementation, may pinpoint yet unknown F^-^ transporters and other molecules in plants specifically targeted by the PGPR to reduce F^-^ ion accumulation in plant tissues, promoting growth under stress. Taken together, this study highlights the promising role of PGPR strains sourced from F^-^-contaminated extreme arid environments in the growth promotion of the model plant *A. thaliana*, leading to the potential development of biofertilizers to mitigate high salt and F^-^ stress under agricultural settings. However, this requires extensive trials on different crop species in the field. The whole genome sequence of *P. megaterium* SI1-IITJ paves the way for the future improvement of this PGPR strain for enhanced ability to mitigate different abiotic stresses in various crops through genome editing and microbiome engineering.

## Declaration of generative AI in scientific writing

The authors state that generative AI was not used to compose or edit the manuscript.

## CRediT authorship contribution statement

Devanshu Verma: Writing – original draft, Investigation. Pinki Sharma: Methodology, Investigation, Writing – original draft. Rishabh Kumar: Investigation, Writing – original draft. Vijesh Prajapat: Investigation. Debankona Marik: Methodology, Investigation. Neelam Jangir: Investigation. Rabisankar Mandi: Investigation. Trisikhi Raychoudhury: Supervision, Resources. Nar Singh Chauhan: Methodology, Investigation, Supervision, Resources. Ayan Sadhukhan: Writing – original draft, Review & editing, Supervision, Funding acquisition, Project administration, Conceptualization.

## Data availability statement

All data are presented within the paper. The transcriptome raw data are submitted in the NCBI Sequence Read Archive with an accession number PRJNA1269765.

## Ethics approval

This study does not require approval from research ethics committees.

## Consent for publication

All authors have checked the draft manuscript and have consented to publication.

## Declaration of competing interest

The authors declare no competing interests.

## Acknowledgments and Funding

AS acknowledges funding from IIT Jodhpur (I/SEED/ASK/20220015) and the Science and Engineering Research Board, Govt. of India (SRG/2022/000169) for financial support. DV obtained a doctoral fellowship from the Department of Biotechnology (DBT), Govt. of India, NJ, and RM from the University Grants Commission (UGC), Govt. of India, and DM from the Ministry of Education, Govt. of India.

## Key message

*Priestia megaterium* SI1-IITJ, root endophyte of Thar Desert plants, mitigates fluoride stress tolerance of *Arabidopsis thaliana* through induction of salt tolerance genes and suppression of transporters and ROS production.

## Notes

### Competing Interest Statement

The authors have declared no competing interest.

