## Supplementary data for "A desert endophyte, *Priestia megaterium* SI1-IITJ, improves fluoride stress tolerance by reducing fluoride content of plant tissues and perturbing salt tolerance and defense genes of *Arabidopsis thaliana*"

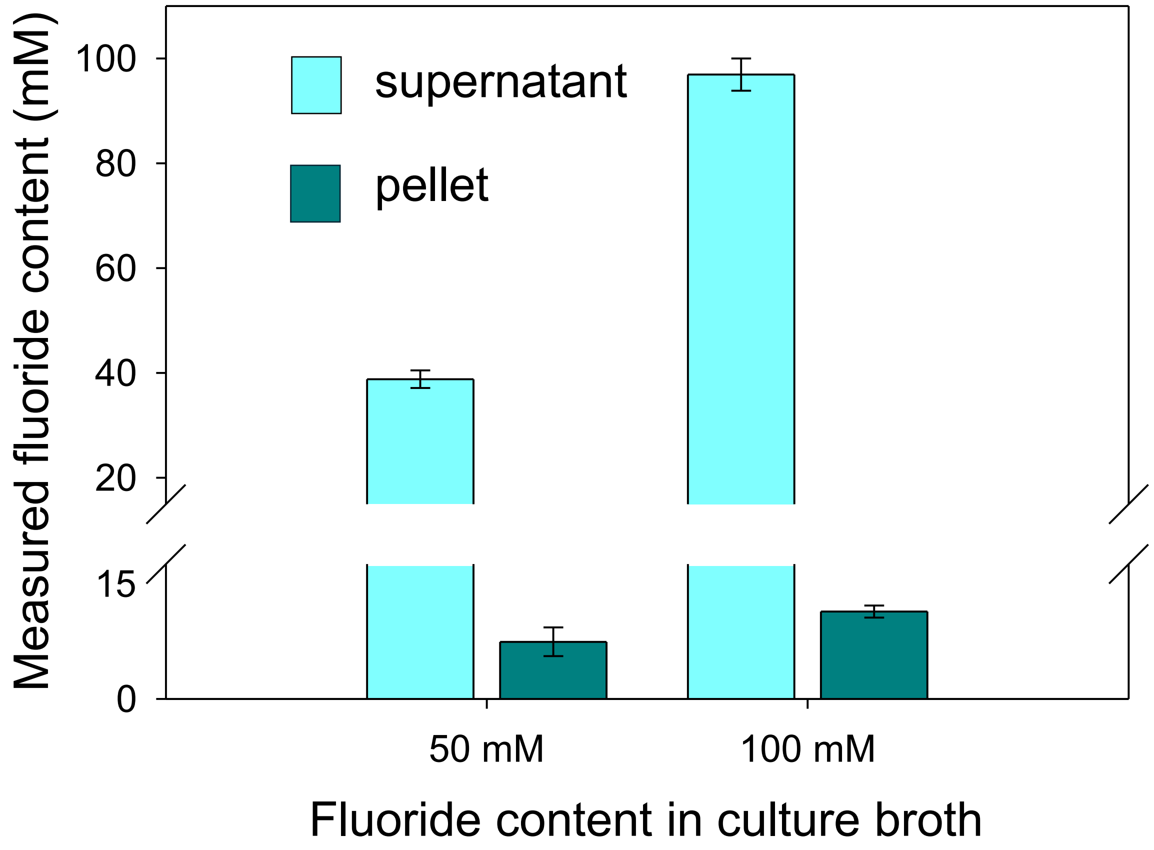


**Fig. S1. Fluoride content in bacteria.** *Priestia megaterium* SI1-IITJ was grown for 24 h in LB broth supplemented with 50 and 100 mM NaF and centrifuged. The estimated fluoride contents in the supernatant and the pellet using a fluoride-sensitive probe are represented (see Methods). Average values of three replicates with standard error are shown.

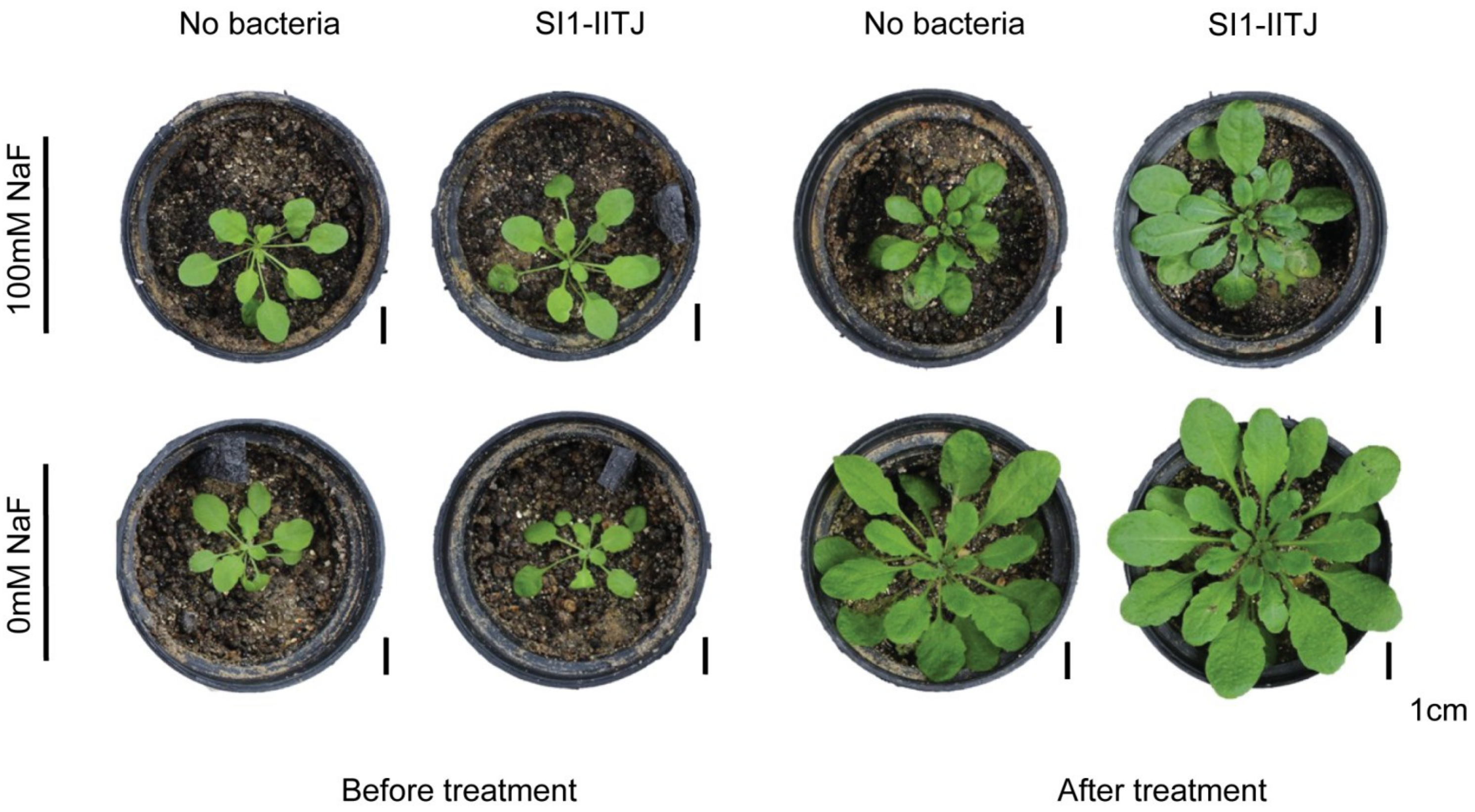


**Fig. S2. Growth promotion of *Arabidopsis thaliana*, ecotype Star-8, by *Priestia megaterium* SI1-IITJ under fluoride stress.** Growth of one-month-old *Arabidopsis thaliana* Star-8 plants in the soil culture experiment is shown. The pots were irrigated with different concentrations of NaF (see Methods). Three biological replicates are shown for each treatment. Black bars represent a scale of 1 cm

**Table S1. Oligonucleotides used in the study**

| Oligonucleotide Name | Sequence (5'->3') |
|---|---|
| 16S Universal Primer_Fw | AGRGTTYGATYMTGGCTCAG |
| 16S Universal Primer_Rv | RGYTACCTTGTTACGACTT |
| AT5G26130_qPCR_Fw | ACTCAAGTTGTGTGGCGAACC |
| AT5G26130_qPCR_Rv | ACGGCCCCTATAATTACCAGGT |
| AT1G19530_qPCR_Fw | AGCAAGAGTAGTGGATGGTGGA |
| AT1G19530_qPCR_Rv | AGTTGTTCGGTCGACCCTCT |
| AT1G12080_qPCR_Fw | GTGGTTGTGGAAGAGGCAGAG |
| AT1G12080_qPCR_Rv | CGGAGTTTCGGTCACTTCCTC |
| AT1G12010_qPCR_Fw | GCGTCGTTTTACAACCCCGG |
| AT1G12010_qPCR_Rv | GCCTGAAACTTGAGTCCGGC |
| AT2G33830_qPCR_Fw | GTAAAACTGTGGCGGCGGTG |
| AT2G33830_qPCR_Rv | GCCCATTCCTCTAGTGGCGA |
| AT3G10020_qPCR_Fw | CCGGCGATGGATGAAGGAGA |
| AT3G10020_qPCR_Rv | CCGGCGATGGATGAAGGAGA |
| *AtUBQ1* _qPCR_Fw | TCGTAAGTACAATCAGGATAAGATG |
| *AtUBQ1* _qPCR_Rv | CACTGAAACAAGAAAAACAAACCCT |

**Table S2 Antibiotic resistance genes of *Priestia megaterium* SI1-IITJ predicted by the Comprehensive Antibiotic Resistance Database (CARD) identifier**

| ARO Term | Detection Criteria | AMR Gene Family | Drug Class | Resistance Mechanism | % Identity of Matching Region | % Length of Reference Sequence |
|---|---|---|---|---|---|---|
| qacJ | protein homolog model | small multidrug resistance (SMR) antibiotic efflux pump | disinfecting agents and antiseptics | antibiotic efflux | 50.48 | 100.93 |
| BcIII | protein homolog model | class A Bacillus cereus Bc beta-lactamase | cephalosporin | antibiotic inactivation | 64.87 | 98.1 |
| vanY gene in vanB cluster | protein homolog model | vanY, glycopeptide resistance gene cluster | glycopeptide antibiotic | antibiotic target alteration | 40.95 | 41.79 |
| qacJ | protein homolog model | small multidrug resistance (SMR) antibiotic efflux pump | disinfecting agents and antiseptics | antibiotic efflux | 37.86 | 96.26 |
| FosBx1 | protein homolog model | fosfomycin thiol transferase | phosphonic acid antibiotic | antibiotic inactivation | 65.22 | 102.9 |
| vanW gene in vanI cluster | protein homolog model | vanW, glycopeptide resistance gene cluster | glycopeptide antibiotic | antibiotic target alteration | 41.33 | 91.69 |
| vanY gene in vanB cluster | protein homolog model | vanY, glycopeptide resistance gene cluster | glycopeptide antibiotic | antibiotic target alteration | 34.21 | 100.75 |
| vanW gene in vanI cluster | protein homolog model | vanW, glycopeptide resistance gene cluster | glycopeptide antibiotic | antibiotic target alteration | 40 | 87.4 |
| qacG | protein homolog model | small multidrug resistance (SMR) antibiotic efflux pump | disinfecting agents and antiseptics | antibiotic efflux | 45.28 | 112.15 |
| vanT gene in vanG cluster | protein homolog model | glycopeptide resistance gene cluster, vanT | glycopeptide antibiotic | antibiotic target alteration | 34.26 | 58.01 |
| vanY gene in vanA cluster | protein homolog model | vanY, glycopeptide resistance gene cluster | glycopeptide antibiotic | antibiotic target alteration | 40.98 | 28.38 |

**Table S3. Mobilome in *Priestia megaterium* SI1-IITJ**

| Mobile Genetic Elements | Details | Position (bp) | Length (bp) | GC Content (%) | Antibiotic Resistance Genes | Drug Class | Drugs | Virulence Gene | Virulence Description |
|---|---|---|---|---|---|---|---|---|---|
| Genomic_island | - | 1573..15831 | 14258 | 32.83 | - | - | - | - | - |
| Prophage | - | 813306..849328 | 36022 | 36.3 | - | - | - | - | - |
| Prophage | - | 62954..69299 | 6345 | 35.11 | - | - | - | - | - |
| Prophage | - | 147921..157678 | 9757 | 38.17 | - | - | - | - | - |
| Prophage | - | 67394..81091 | 13697 | 38.38 | - | - | - | - | - |
| Integrative_Conjugative_Elements | - | 17915..61561 | 43646 | 33.24 | - | - | - | - | - |
| IS/Tn | ISBce13 | 7328..8206 | 878 | 39.75 | - | - | - | - | - |
| IS/Tn | IS240C | 103353..103874 | 521 | 36.08 | - | - | - | - | - |
| IS/Tn | ISBce13 | 130235..131113 | 878 | 39.64 | - | - | - | - | - |
| IS/Tn | ISBsu1 | 70459..71313 | 854 | 39.81 | - | - | - | - | - |
| IS/Tn | ISBsu1 | 84101..84955 | 854 | 40.28 | - | - | - | - | - |
| IS/Tn | ISBsu1 | 93508..93678 | 170 | 41.76 | - | - | - | - | - |
| IS/Tn | ISBce13 | 68669..69547 | 878 | 39.86 | - | - | - | - | - |
| IS/Tn | ISBwe3 | 103752..104447 | 695 | 37.55 | - | - | - | - | - |
| IS/Tn | ISBwe2 | 56709..56867 | 158 | 39.24 | - | - | - | - | - |
| IS/Tn | ISBwe2 | 74747..75025 | 278 | 39.57 | - | - | - | - | - |
| IS/Tn | ISBth6 | 76448..76753 | 305 | 36.07 | - | - | - | - | - |
| IS/Tn | ISBsu1 | 30372..30794 | 422 | 39.81 | - | - | - | - | - |
| IS/Tn | ISBame1 | 15107..15517 | 410 | 40 | - | - | - | - | - |
| IS/Tn | ISBce11 | 50260..51690 | 1430 | 39.72 | - | - | - | - | - |
| IS/Tn | ISBwe3 | 62892..63602 | 710 | 39.44 | - | - | - | - | - |

Mobile genetic elements are shown along with their respective position (in base pairs) in the genome of *Priestia megaterium* SI1-IITJ. The mobile elements were detected by the VRprofile2 tool version 2.0.
